# Conformational dynamics of the membrane-anchored foldase LipH from *Pseudomonas aeruginosa* governs recognition and release of its client lipase

**DOI:** 10.64898/2026.03.12.711337

**Authors:** Max Busch, Jennifer Loschwitz, Athanasios Papadopoulos, Jens Reiners, Wieland Steinchen, Vincenzo Calvagna, Sander H.J. Smits, Karl-Erich Jaeger, Alexej Kedrov

**Affiliations:** Heinrich Heine University Düsseldorf, Synthetic Membrane Systems, Düsseldorf, Germany; Heinrich Heine University Düsseldorf, Center for Structural Studies, Düsseldorf, Germany; Marburg University, Center for Synthetic Microbiology & Dept. Chemistry, Marburg, Germany; Heinrich Heine University Düsseldorf, Structural Biochemistry, Institute of Biochemistry, Düsseldorf, Germany; Heinrich Heine University Düsseldorf, Interfaculty Center for Membrane Research, Düsseldorf, Germany; Heinrich Heine University Düsseldorf, Institute of Molecular Enzyme Technology, Düsseldorf, Germany; Forschungszentrum Jülich GmbH, Institute of Bio- and Geosciences IBG-1: Biotechnology, Jülich, Germany

## Abstract

The lipase LipA from *Pseudomonas aeruginosa* is an extracellular enzyme that plays an important role in bacterial infections. Prior its export via the type II secretion system, LipA requires the cognate membrane-anchored foldase LipH for maturation in the periplasm. Though structural studies elucidated the architecture of the LipH:lipase complex, how the full-length, membrane-tethered foldase recognizes, folds, and releases its client has remained barely understood. Here, we combine *in silico* and *in vitro* analysis to resolve the conformational dynamics and function of the full-length LipH in a membrane context. Simulations reveal that the membrane-anchored LipH is highly dynamic, sampling a broad ensemble of conformations. The chaperoning cavity is transiently closed or occluded by both the proximal membrane and the linker polypeptide, which is further confirmed by structural analysis. Despite the steric hindrance, the full-length LipH reconstituted into lipid-based nanodiscs and amphipols efficiently activates LipA, though it displays substantially reduced affinity for the client. We propose that the negatively charged membrane promotes release of the folded client, enabling multiple chaperoning cycles. Hydrogen/deuterium exchange analysis reveals that the MD2 domain of LipH in engaged in stable interactions with the lipase, whereas MD1 contacts are transient. Consistently, LipH readily captures N-terminal fragments of LipA, indicating that initial recognition relies on local interactions via the MD2 domain. Together, our results show how membrane coupling and intrinsic conformational plasticity modulate the function of the steric chaperone, and suggest that the membrane-anchored LipH balances capture, folding, and release of the client LipA to enable its efficient secretion.

## Introduction

Protein secretion in bacteria plays a central role in colonization, invasion and host-pathogen interactions, as, among other clients, it enables transport of virulence factors across cellular membranes. In pathogenic bacteria, such as an opportunistic human pathogen *Pseudomonas aeruginosa*, specialized secretion systems facilitate the delivery of toxins and effector proteins, either into the environment or into host cells ^1^. The type II secretion system (T2SS), also referred as the general secretion pathway due to its high conservation among Gram-negative bacteria and broad specificity, mediates export of folded client proteins from the periplasm into the extracellular space ^2^. In the opportunistic pathogen *Pseudomonas aeruginosa*, the lipase A (LipA) is a highly abundant secretory protein, which is exported via Xcp-T2SS ^3,4^. Due to its association with biofilms and contribution to lysis of host cells, LipA is considered as a virulence factor ^4^, and understanding the route of LipA folding and secretion would be valuable for combating the pathogenesis of *P. aeruginosa*. Biogenesis and secretion of the lipase occur in distinct steps, starting with translation of the precursor protein and its translocation into the periplasm via the SecYEG:SecA route (Figure 1A). There, maturation of LipA, i.e. cleavage of the signal sequence, takes place, followed by folding into its enzymatically active form, recognition by T2SS and secretion across the outer membrane ^5^. Notably, LipA secretion is essentially dependent on the specific chaperone LipH, also referred as lipase-associated foldase or *Lif* ^5,6^. Encoded in one operon with LipA, LipH is a membrane-anchored chaperone that recognizes its cognate client in the periplasm and facilitates its correct folding, so the lipase overcomes the energetic barrier towards the catalytically active conformation ^5,7-9^. In a following step, the folded LipA should be released from LipH and handed over to the dedicated Xcp-T2SS, potentially via interactions with the XcpP protein in *P. aeruginosa* ^10^, and then it is expelled upon a piston-like movement of the T2SS pseudopilus ^4^.

**Figure 1.**
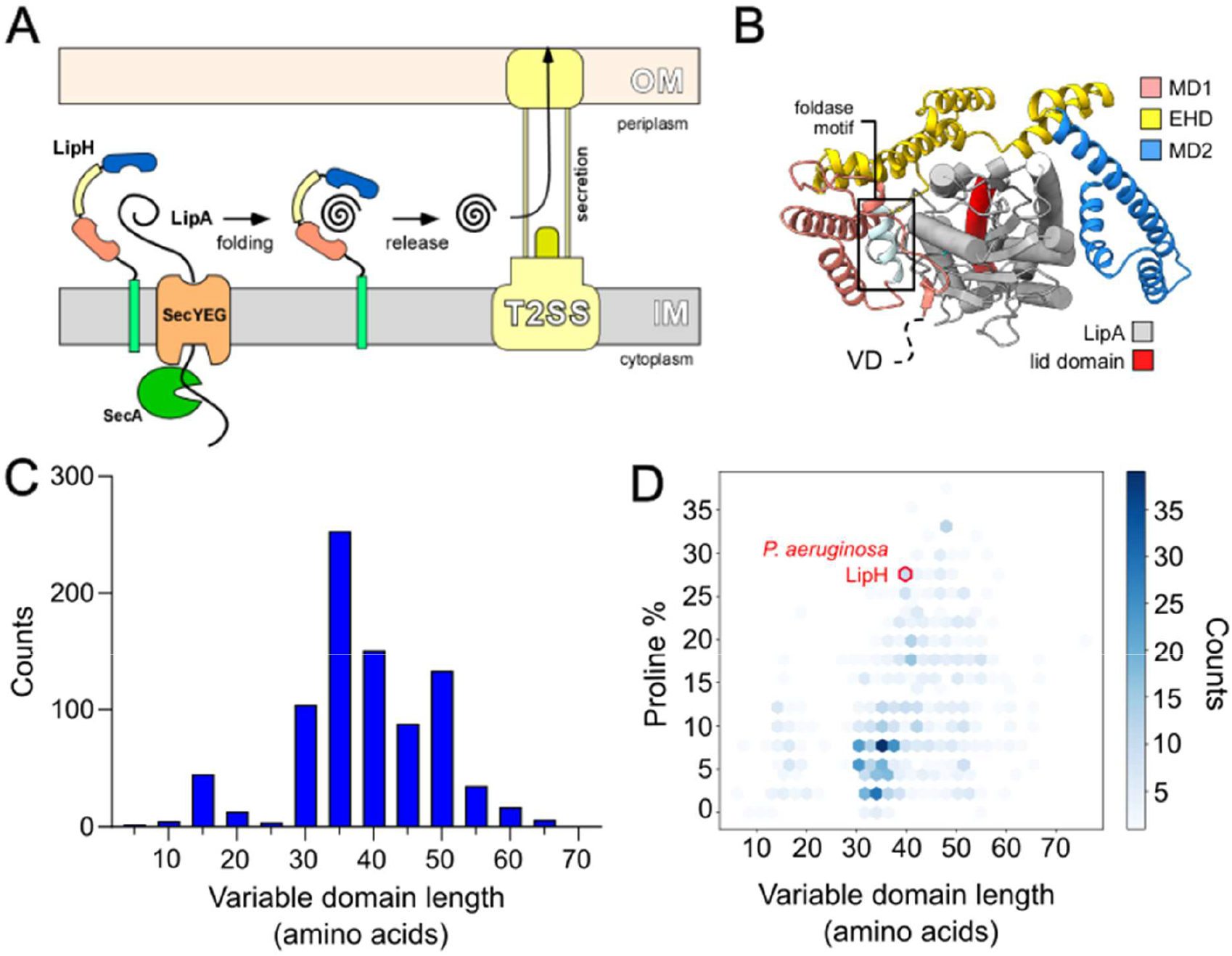
Architecture of the lipase-specific foldase LipH. **(A)** Scheme of LipA secretion route in *P. aeruginosa*. Translocation across the inner membrane (IM) via the SecYEG:SecA machinery is followed by recognition via the membrane-anchored steric chaperone LipH. LipH facilitates folding and activation of the lipase. After being released, LipA is competent for secretion via the type II secretion system (T2SS). **(B)** AlphaFold3-based model of *P. aeruginosa* LipH:LipA complex. The chaperoning domain of LipH (residues 63-340) is shown as a helical ribbon, with the structural elements color-coded. The N-terminal fragment (residues 1-62), including the variable domain (VD) is omitted for clarity. **(C)** Histogram of the variable domain length distribution among the identified lipase-specific foldases (in total, 858 sequences). **(D)** Distribution of the polypeptide length and the proline content of the variable domains among the identified lipase-specific foldases. Position of *P. aeruginosa* LipH^VD^ is highlighted in red.

The lipase-foldase system is not limited to *Pseudomonas*, but also found in the genera *Burkholderia, Acinetobacter*, and *Vibrio* ^5,11^. The crystallized foldase:lipase complexes from *Burkholderia glumae* and *Acinetobacter baumannii* reveal nearly identical structures ^12,13^, and the earlier model of *P. aeruginosa* LipH:LipA ^14^, as well as the AlphaFold-based prediction manifest a highly similar organization (Figure 1B, Suppl. Figure 1). Briefly, the chaperoning domain of LipH is helically folded and it consists of two mini-domains, MD1 and MD2, connected via an extended helical domain (EHD), forming together a crescent-like structure, also called “headphone” shape. MD1 contains the characteristic sequence R_94_NLFDYFLSA_103_ found among the foldases (consensus motif RXXFDY(F/C)L(S/T)A, where X can be any amino acid) ^5^. The available structures and the AlphaFold model reveal a large interaction interface between the chaperone and the lipase. Key contact points within the *P. aeruginosa* LipA:LipH complex include helix 1 of LipH MD1, where Ser-112 is positioned near Gln-275 of the lipase; helix 5 (part of the EHD), where Arg-199 may interact with Asn-26 of LipA; and helix 11 (part of MD2), where Arg-280 and Arg-327 potentially build salt bridges with LipA Glu-66 and Glu-58, respectively (the numbering corresponds to *P. aeruginosa* LipA without the signal sequence). Notably, the complex assembly does not involve the helix 5 of the lipase, also referred as a lid domain, that regulates access to the lipase catalytic site formed by residues Ser-82, Asp-229 and His-251 ^14^. The available structures, as well as the AlphaFold models, show a displacement of the lid domain (Suppl. Figure 2), and a fatty acid can be docked via AlphaFold within the open conformation of LipA, while still bound to LipH, in agreement with functional studies, where LipA is enzymatically active also when bound to LipH ^15,16^.

Extensive functional and structural studies on LipH:LipA interactions have commonly focused on the chaperoning domain of LipH, leaving the preceding N-terminal region out of consideration, although it constitutes nearly 20 % of the full-length LipH in *P. aeruginosa*. The region consists of an N-terminal transmembrane helix (TMH) followed by a non-conserved linker, referred as variable domain (VD). The region serves as a tether and prevents LipH co-secretion with the client lipase ^5^, so the membrane-anchored foldase may sequentially facilitate the fold and release of multiple LipA molecules. The VD linker of *P. aeruginosa* LipH is 41 residues long, of which eleven are prolines, and the AlphaFold-based model suggested lack of secondary structure in the region. The composition of the linker in *P. aeruginosa* LipH stands out among other lipase-related foldases: Analysis of 858 LipH homologs suggested that the linker length commonly occurs within 30-55 residues (Figure 1C), but the proline content is typically below 10 %, while it reaches 27% for *P. aeruginosa* LipH (Figure 1D). Such high proline content is often observed in disordered polypeptide chains ^17^, which may enable movements of the chaperoning domain above the membrane to escape the steric hindrance, as it would be required for capturing the client lipase and/or for downstream interactions with T2SS ^7,18^. However, no experimental studies addressing the dynamics and interactions of the full-length LipH are available, despite their relevance to folding and maturation processes of the client virulence factor.

Here, we combine molecular dynamics simulations, structural characterization, and biochemical analysis to probe the conformational dynamics of *P. aeruginosa* LipH, its interaction with the client lipase, and the effect of the membrane environment on the chaperoning activity. We show that LipH is highly flexible due to the unstructured linker VD and the intrinsically dynamic chaperoning domain, enabling it to sample a large periplasmic space. We propose that the client LipA is initially recognized at its N-terminal fragment through interactions with the MD2 domain of LipH. Once the full-length LipA binds, the loosely bound LipH MD1 domain likely contributes to positioning, folding and eventual release of the folded lipase, which is assisted by the proximate membrane environment. Together, our work proposes a mechanistic model for the steric foldase function related to the pathogenicity of *P. aeruginosa*.

## Results

### Dynamics of the full-length LipH at the membrane interface

Aiming to resolve the dynamic interactions of LipH, we first set out to characterize the full-length protein (“LipH^FL^”: Suppl. Figure 3) at a membrane interface via all-atomistic molecular dynamics (MD) simulations (Figure 2A). LipH^FL^ was anchored with its N-terminal helix in the phospholipid bilayer composed of phosphatidylethanolamine (PE, 75 mol %), phosphatidylglycerol (PG, 20 mol %) and cardiolipin (CL, 5 mol %), thus mimicking the bacterial inner membrane ^19^. The simulation box of 22 x 22 x 19 nm^3^ was designed to be sufficiently large to trace the movements of LipH domains above the membrane interface, and the non-restricted atomistic simulations allowed to reconstruct the dynamics of the chaperoning domain, as well as the VD linker. Furthermore, two different environments were studied, containing NaCl of either 25 mM or 150 mM concentration. The low ionic strength approximated the periplasmic conditions of *P. aeruginosa* in the soil habitat, and the high ionic strength reflected the environment upon the tissue or lung invasion ^20^.

**Figure 2.**
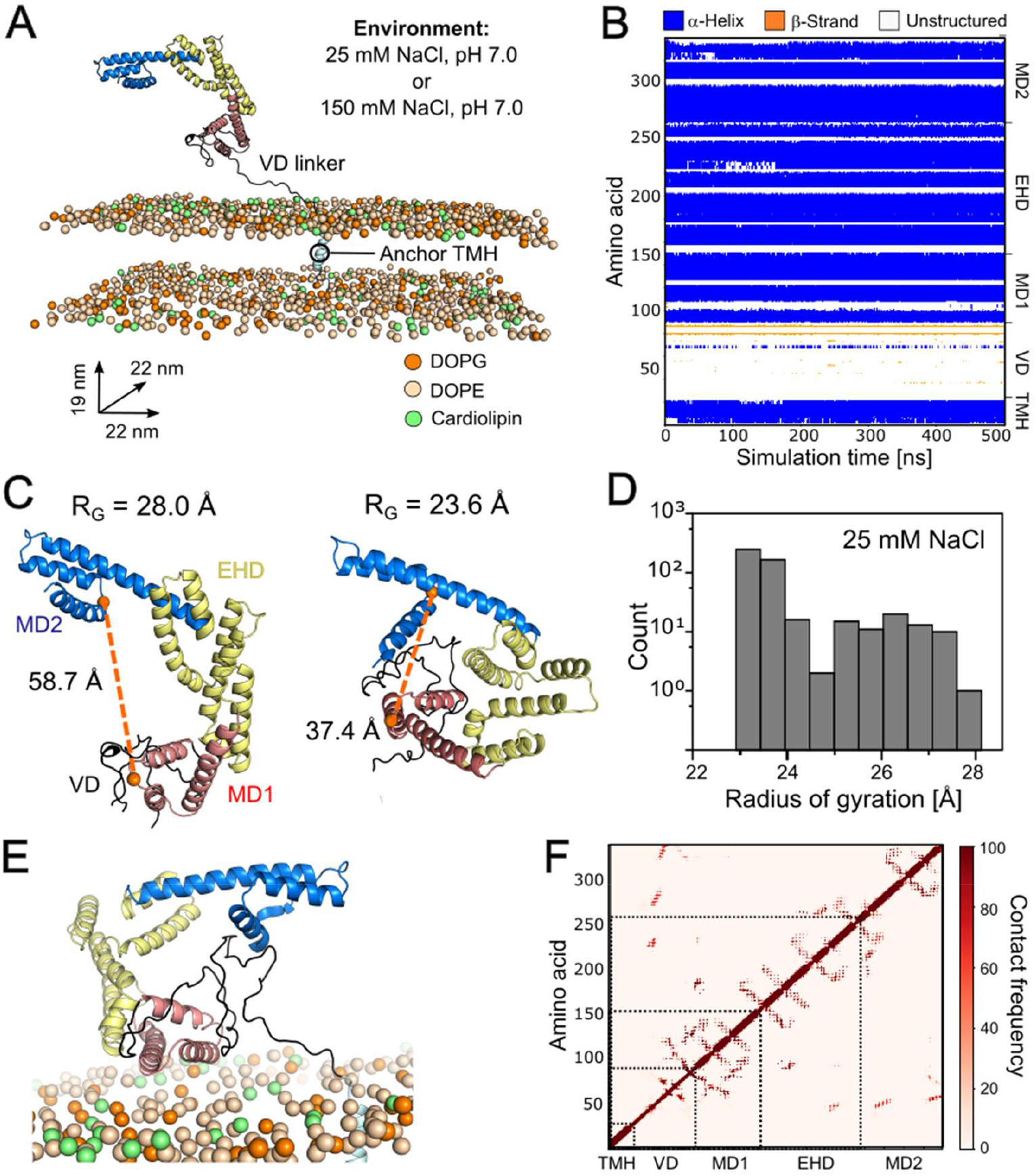
Molecular dynamics simulation of the full-length LipH at the lipid membrane interface. **(A)** Design of the molecular dynamics simulation system. LipH^FL^ was anchored with its N-terminal helix (TMH, cyan) in a native-like lipid membrane. The dimensions of the designed simulation box are indicated. **(B)** The color-coded secondary structure of LipH^FL^ along a simulation course (25 mM NaCl) is plotted against the simulation time. **(C)** Dynamics of the chaperoning domain within LipH^FL^. Left: Conformation at the beginning of the simulation. Right: An exemplary conformation of a compact state acquired along the simulation. To analyse the dimensions of the domain, the distance between the Cα atoms of Glu-107 (MD1) and Glu-321 (MD2) was monitored (indicated as the orange spheres and the dashed line). **(D)** Distribution of the radius of gyration of the LipH chaperoning domain along a simulation course (25 mM NaCl). **(E)** A representative conformation of LipH^FL^ in proximity to the lipid membrane shows extensive contacts of the unstructured VD polypeptide within the chaperoning domain. **(F)** Map of intramolecular contacts within LipH^FL^ along a simulation course indicates interactions of the VD linker with MD1, EHD and MD2 domains.

During all performed simulations, LipH^FL^ manifested little to no change in the secondary structure content, i.e. the chaperoning domain and the membrane anchor retained their α-helical fold, and the VD polypeptide remained largely unstructured, with transient formation of short β-strands (Figure 2B and Suppl. Figure 4). However, the architecture of the chaperoning domain was not rigid, as the mini-domains MD1 and MD2 underwent extensive movements, causing transient widening and closing of the chaperoning cavity. Thus, the distance between the mini-domains measured for the chosen residues Glu-107 (MD1) and Glu-321 (MD2) varied from 32.93 to 68.12 Å, and the hydrodynamic radius of the chaperoning domain ranged from 23.07 to 29.77 Å in 25 mM NaCl and from 23.13 to 29.17 Å in 150 mM NaCl (Figure 2C and D, Suppl. Figures 5 and 6). At the end of the simulations, the chaperoning domain commonly acquired more compact states, both at the low and high ionic strength. Notably, multiple conformations of LipH showed the VD polypeptide ingressing the chaperoning cavity, so the MD1 domain, and to a lesser extent, MD2 repeatedly interacted with the unstructured region (Figure 2E and F, Suppl. Figure 7). Together with the movements of the MD1/MD2 domains, this unexpected occlusion could sterically hinder specific LipH:LipA interactions and, therefore, affect folding and activation of the client lipase.

The interactions of LipH with the membrane, first of all the electrostatic interactions with the lipid head groups, must be decisive for the conformational freedom of the chaperone domain and its accessibility for the client. We noted that the charged amino acids are distributed in a non-uniform manner over LipH: While the cationic residues are predominantly found within the LipA binding interface, the anionic residues are exposed at the outer surface, with a pronounced cluster within MD1 (Figure 3A). This negatively charged cluster should ensure electrostatic repulsion from the abundant anionic lipids, such as PG and CL, at the membrane. Indeed, when only the chaperoning domain (“LipH^Chap^”, Suppl. Figure 3) was simulated in absence of the TMH anchor and the VD polypeptide, it was found exclusively in the solvent, forming no contacts with the lipid leaflet (Figure 3B and Suppl. Figure 8). In contrast, when examining the dynamics of LipH^FL^, we observed multiple events, when the minimal protein-lipid distance measured for the chaperoning domain approached zero, reflecting a close contact with the membrane (Figure 3B and Suppl. Figure 9). When comparing different ionic strengths, LipH docking at the membrane interface was observed in a single simulation at 25 mM NaCl, where both MD1 and MD2 domains formed a long-lasting contact with the lipid leaflet, thus blocking the access to the chaperoning cavity (Figure 3 B-E; Suppl. Figure 9). Two other simulations at the low ionic strength showed LipH exposed to the solvent, being accessible for the client binding. In contrast, all three simulations at 150 mM NaCl indicated LipH:membrane interactions (Figure 3E and Suppl. Figure 8B and C). Here, the negatively charged MD1 repeatedly approached the surface, so the electrostatic repulsion was apparently compensated by the elevated salt concentration. Thus, LipH:LipA interactions may be hindered by the membrane interface at the elevated salt concentrations, and efficient secretion of LipA would require accessory holdase chaperones ^16^.

**Figure 3.**
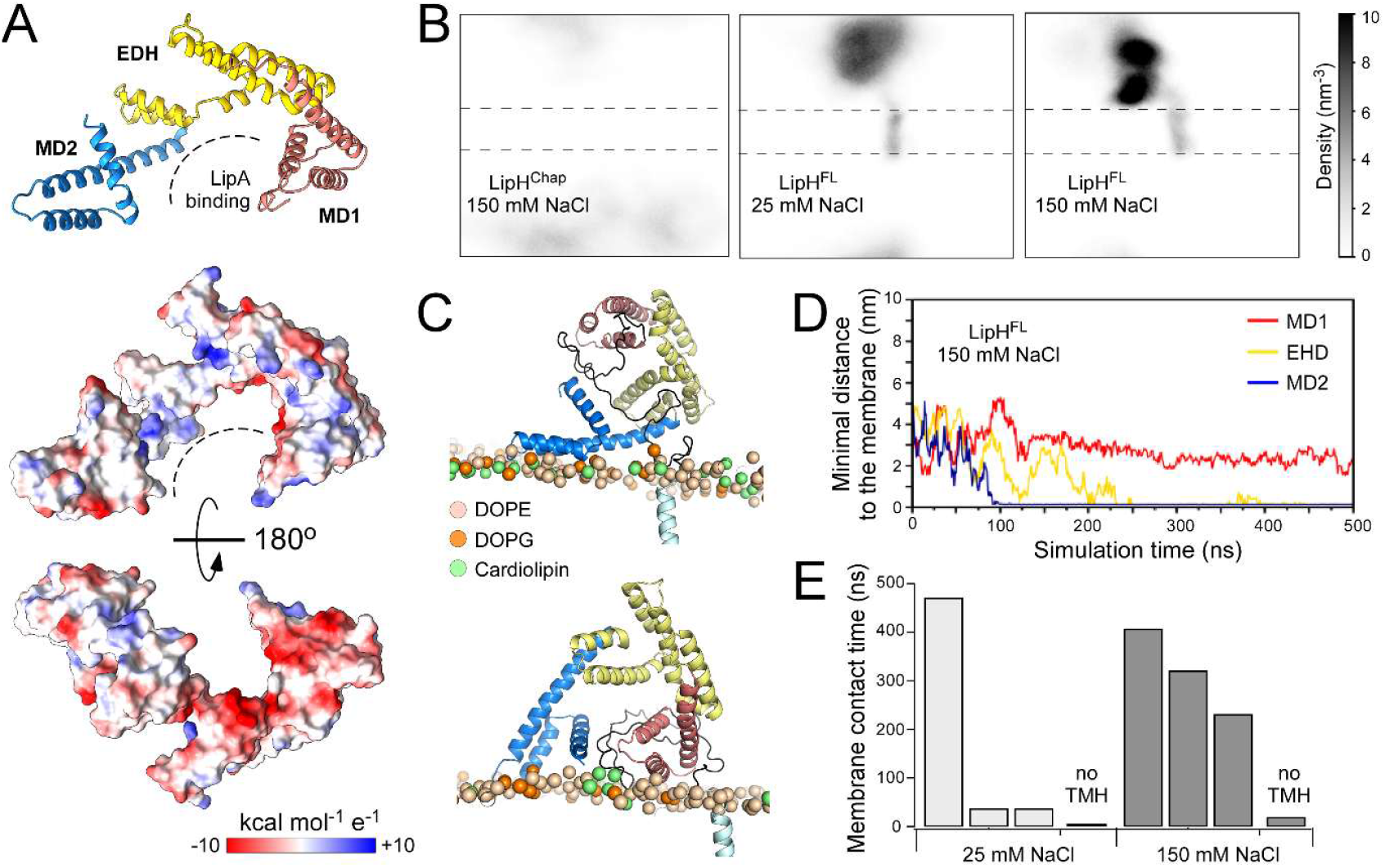
LipH interactions with the lipid membrane interface. **(A)** AlphaFold-based model of *P. aeruginosa* LipH suggests presence of a negatively charged cluster of residues within the MD1 domain. Above: A ribbon visualization of the LipH chaperoning domain; below: the corresponding molecular surface coloured according to the local electrostatic potentials (blue = cationic; red = anionic; the scale bar shown below). **(B)** Localization of the soluble chaperoning domain LipH^Chap^ and the lipid-anchored LipH^FL^ within the simulation box shown as density maps. The lipid bilayer borders are indicated as dashed lines. **(C)** Examples of conformations acquired by LipH^FL^ upon contacting the membrane interface. **(D)** Minimal distances of the individual LipH domains to the membrane over the course of a simulation. **(E)** Duration of LipH:membrane contacts in individual simulations at 25 and 150 mM NaCl.

### Dynamics of the LipH chaperoning domain

The MD simulations of the membrane-anchored LipH^FL^ suggested extensive dynamics within the chaperone in the absence of the client. Previously, single-molecule FRET studies and MD simulations on the isolated LipH^Chap^ construct have led to the qualitatively similar conclusions about the movements of the constituting domains ^18^. However, the N-terminal VD linker was omitted from consideration, so its putative interactions with the chaperoning domain have not been experimentally tested. To do so, we designed a soluble LipH variant consisting of the chaperoning domain with the preceding VD polypeptide (“LipH^VD^”, Suppl. Figure 3) and carried out its biochemical and structural analysis (Figure 4A). The purified protein was monomeric as tested by size-exclusion chromatography coupled to multi-angle light scattering (SEC-MALS) (Figure 4B),and small-angle X-ray scattering (SAXS) was further performed to assess its conformational dynamics. SAXS recordings confirmed the monomeric state of LipH^VD^ (Suppl. Table 1) but also provided estimates of its spatial dimensions and the conformational heterogeneity in the molecular ensemble. The pair-distance distribution function *p(r)*, i.e. the distance between each individual pair of atoms of the molecule, suggested the maximal dimensions of the molecule (r_*max*_) of 12.61 nm a radius of gyration (*R*_*g*_) of 3.41 nm (Figure 4C and Suppl. Figure 10B). The broad width of Kratky plot reflected the typical behavior of a flexible and elongated molecule, in line with the *p*(r) function (Suppl. Figure 10C). To investigate the flexibility of LipH^VD^ and to assess its possible conformations based on SAXS data, we employed the Ensemble Optimization Method (EOM), assuming independent movements of the individual sub-domains MD1, EHD and MD2, and the N-terminal VD polypeptide. The resulting *R*_*g*_ distribution of the molecular ensemble exhibited a bimodal profile with maxima at 2.8 nm and 3.8 nm (*χ*^2^ value of 1.026, CorMap *P*-value of 0.553) (Suppl. Figure 10D and E). This indicated a distinct heterogeneity among LipH^VD^, and the bimodal distribution could be described well with structural models containing the VD polypeptide within the chaperoning domain (the compact state, 60 % of the conformational ensemble) or stretched away from the protein (the extended state) (Figure 4E and Suppl. Figure 10F), in line with the MD simulations. Furthermore, the SAXS data and the EOM-based models also implied substantial movements of the sub-domains MD1, EHD and MD2, thus experimentally confirming the extensive flexibility of the chaperone.

**Figure 4.**
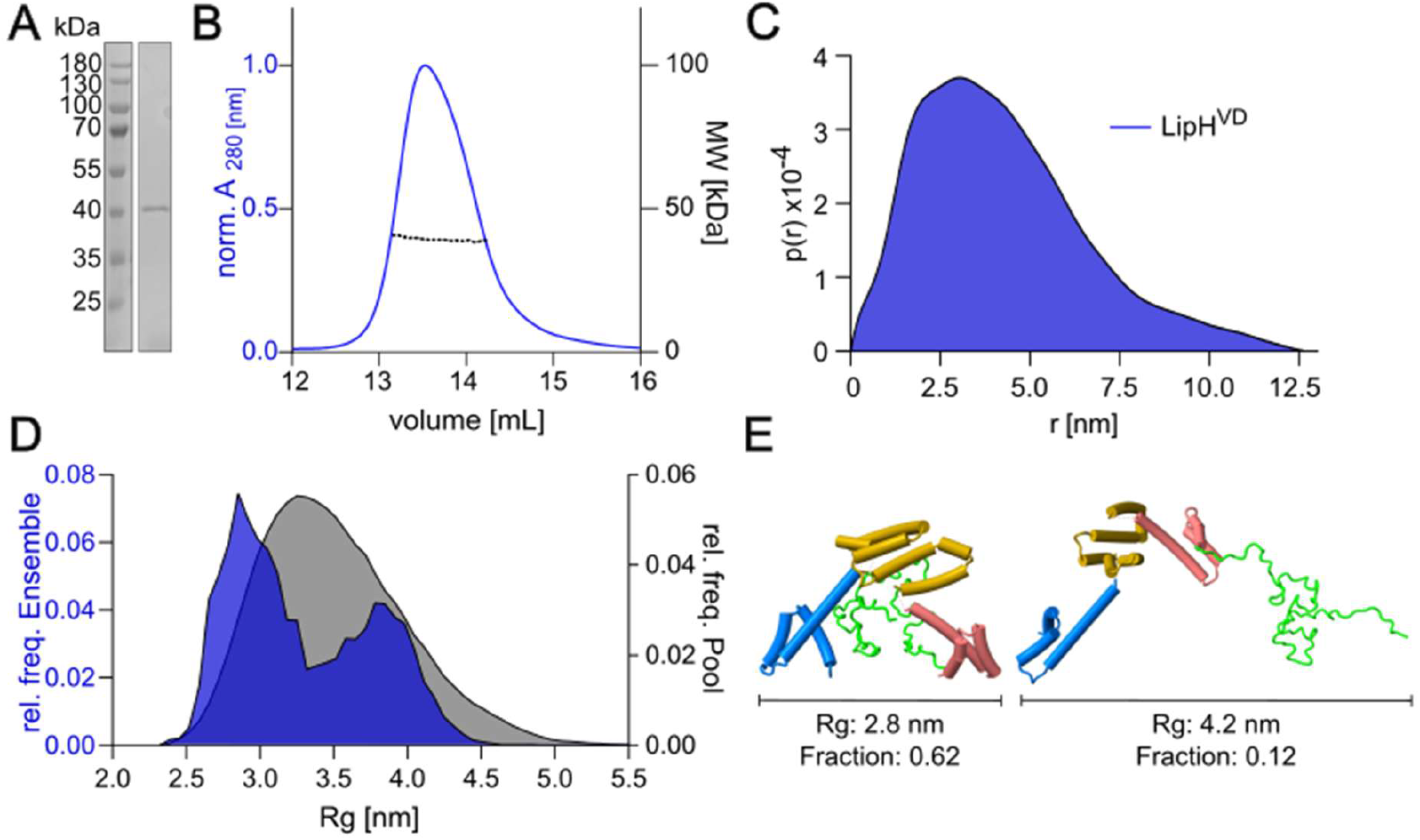
Conformational dynamics of LipH *in vitro*. **(A)** SDS-PAGE of the isolated soluble LipH^VD^. **(B)** SEC-MALS chromatograms of LipH^VD^. Blue line shows the normalized protein absorbance at 280 nm (left Y-axis). Dashed line indicates the molecular weights in kDa (right Y-axis). **(C)** The pair distance distribution function *p(r)* of LipH^VD^ as determined by SAXS. **(D)** Gyration radii distribution (*R*_*g*_) of LipH^VD^ as determined by SAXS. The distribution of the pool models used for fitting is shown in grey; the ensemble of selected EOM models is shown in blue. **(E)** Representative AlphaFold3-based models of LipH^VD^ used for EOM-fitting the SAXS data. The remodeled parts of the N-terminal polypeptide are shown in green loops. Gyration radii *R*_*g*_ and the occupancy of each fraction in the EOM pool are indicated. Further models are shown in Suppl. Figure 10.

**Table 1.**
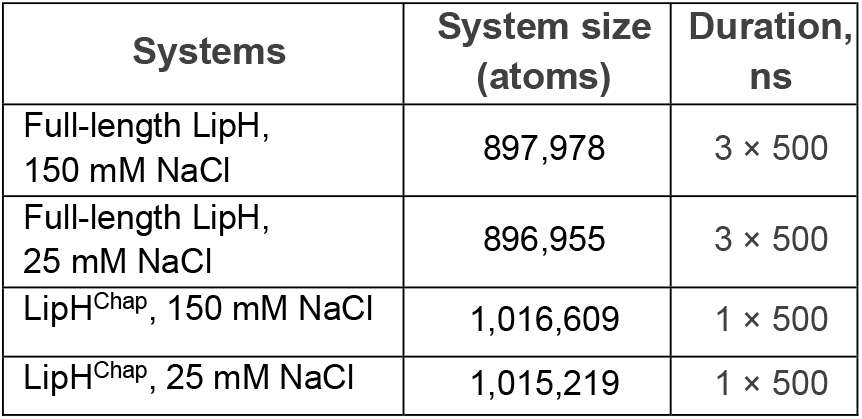
The list of the performed simulations.

| Systems | System size (atoms) | Duration, ns |
| --- | --- | --- |
| Full-length LipH, 150 mM NaCl | 897,978 | 3 × 500 |
| Full-length LipH, 25 mM NaCl | 896,955 | 3 × 500 |
| LipH<sup>Chap</sup>, 150 mM NaCl | 1,016,609 | 1 × 500 |
| LipH<sup>Chap</sup>, 25 mM NaCl | 1,015,219 | 1 × 500 |

### Biochemical characterization of the full-length LipH

MD simulations reveal that both the steric hindrance and electrostatic interactions with the membrane affect the conformation of LipH^FL^, and so they support the idea that the proximate lipid bilayer influences LipH:LipA interactions. To scrutinize the effect of the membrane on the chaperoning properties, we set out to isolate the full-length foldase for the first time and study its interactions with LipA after reconstitution into membrane mimetics, also in comparison with the soluble construct LipH^VD^. The N-terminally tagged LipH^FL^ was heterologous expressed in *E. coli*, extracted with help of non-ionic detergents, DM,DDM, Cymal-6 or LMNG, and isolated via IMAC and subsequent SEC (Figure 5A). With the calculated molecular mass of 39 kDa, the recombinant LipH^FL^ appeared as a band at ∼43 kDa, so it demonstrated a slower migration upon SDS-PAGE (Figure 5B), which is a common behaviour of single-spanning membrane proteins ^21^. While each LipH^FL^ isolation resulted in a sharp peak upon SEC, the elution volume was detergent-specific, varying from 10.5 to 12.5 mL (Figure 5A). To examine whether the shift was caused by differences in LipH^FL^ oligomeric state or by the dimensions of the detergent micelles, we performed SEC-MALS in DDM and Cymal-6, two detergents will well-characterized micellar weights (70 kDa for DDM, 30 kDa for Cymal-6) ^22^. The protein mass calculated for both samples was in the range of 70-80 kDa (Suppl. Figure 11), indicating that LipH^FL^ was purified as a dimer. The dimerization was attributed to the membrane-anchoring helix, since the LipH^VD^ construct remained monomeric even at the elevated concentrations employed in SAXS measurements.

**Figure 5.**
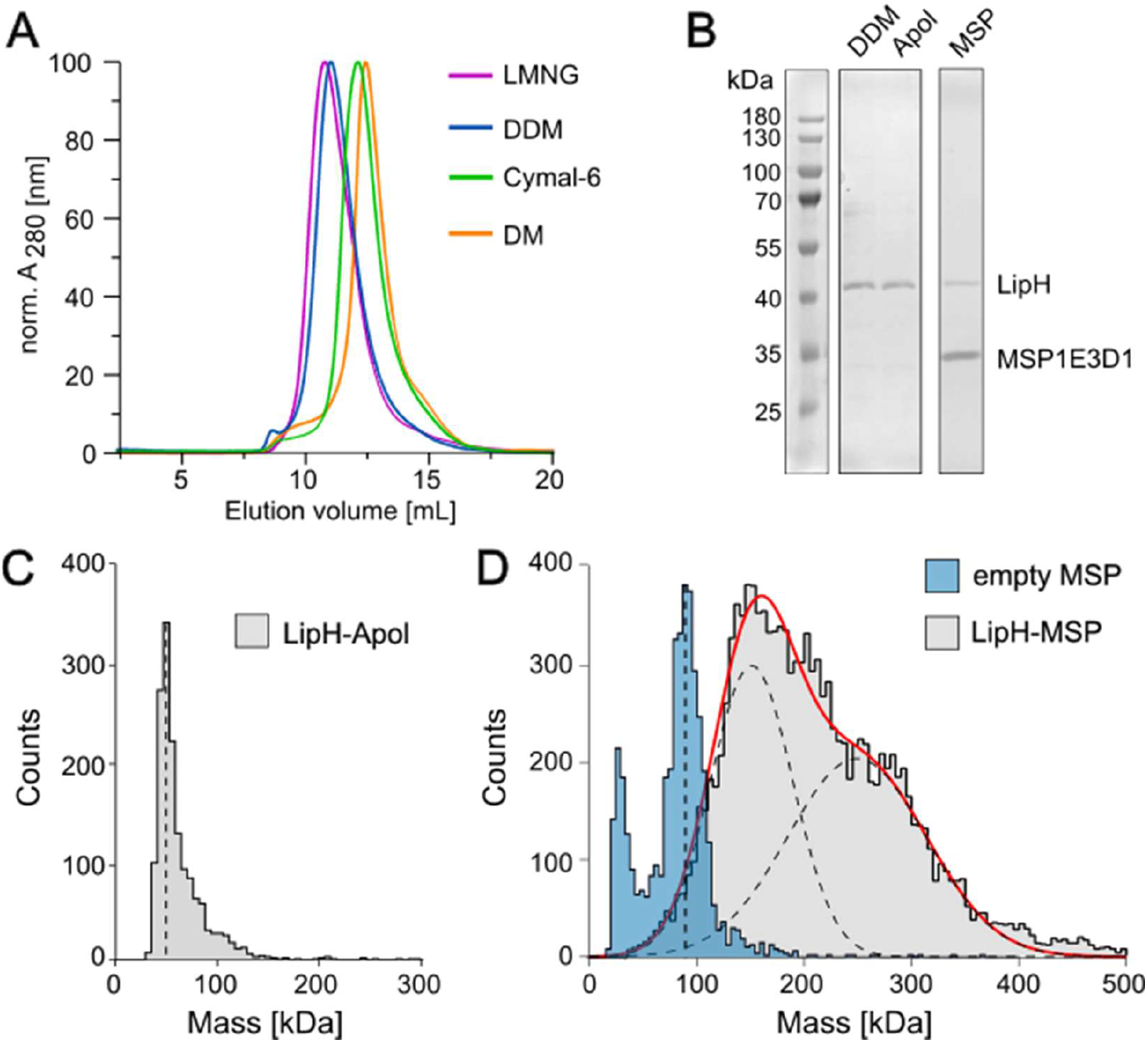
Isolation and characterization of the full-length LipH. **(A)** Size-exclusion chromatography profiles of LipH^FL^ purified in the indicated detergents. **(B)** SDS-PAGE of LipH^FL^in the detergent micelles (“DDM”), amphipols (“Apol”), and nanodiscs (“MSP”). **(C)** Mass distribution of the amphipol-reconstituted LipH (“LipH-Apol”) determined by single-molecule mass photometry. The dashed line indicates the distribution peak at approx. 50 kDa. **(D)** Mass distribution of the MSP1E3D1-based particles, with reconstituted LipH^FL^ (“LipH-MSP”) and without it (“empty MSP”). For the empty MSP, the dashed line indicates the distribution peak at approx. 90 kDa. Fitting the LipH-MSP distribution with two Gaussian peaks (dashed lines) suggests presence of two populations with mean masses of approx. 150 kDa and 250 kDa.

The DDM-solubilized LipH^FL^ was further used for reconstitution in membrane mimetics, i.e. polymer-based particles and lipid-based nanodiscs ^23,24^. Upon removal of the detergent, the amphipathic polymer (amphipol) A8-35 wraps over the hydrophobic surfaces of the target membrane protein ^23^, and the solvent-exposed carboxylate groups ensure solubility of the assembled particle, also in absence of additional lipids. Differently, the nanodiscs consist of a lipid bilayer with an embedded target membrane protein, and the assembly is stabilized by a dimer of the membrane scaffold protein (MSP) bound at the periphery. The MSP1E3D1 scaffold employed here produces nanodiscs of ∼12 nm diameter ^25^, which were loaded with DOPC:DOPG lipids (70:30 molar ratio). LipH^FL^ reconstituted in amphipols (“LipH-Apol”) and nanodiscs (“LipH-MSP”) were analysed by single-molecule mass photometry to determine the molecular weight of the formed particles, and to resolve the oligomeric state of the chaperone. The amphipol-based sample manifested a sharp peak at 50 kDa (Figure 5C), so LipH^FL^ was embedded as a monomer (40 kDa), and the excess mass could be assigned to the polymer ^26^. Differently, LipH-MSP showed a broad mass distribution reflecting the heterogeneity within the ensemble (Figure 5D). The distribution peaked at 150 kDa, and its tail could be fitted with a second Gaussian centred at 250 kDa. As the lipid-loaded nanodiscs in absence of LipH^FL^ (“empty MSP”) manifested a narrow distribution around 90 kDa, the excess masses measured for LipH-MSP likely corresponded to dimers of LipH^FL^ (peak at 150 kDa) and higher oligomers (peak at 250 kDa). Thus, the nanodiscs captured multiple copies of LipH^FL^, possibly in alternating topologies. However, the individual chaperoning domains should be exposed to the solvent being accessible for interaction with the client LipA.

### Full-length LipH may conduct multiple chaperoning cycles

Previous studies showed that the isolated chaperoning domain LipH^Chap^ prevents aggregation of the urea-denatured lipase LipA and facilitates its correct folding and activation *in vitro* ^27 16^. With the assembled set of LipH variants, i.e. LipH^VD^, LipH^FL^-Apol and LipH^FL^-MSP, we set out to analyse the functionality of the chaperone and to identify potential effects of the N-terminal domain and the membrane environment. For example, both occlusion of the binding pocket by the VD polypeptide, as seen in SAXS and MD simulations, and interactions with the membrane surface may suppress the recognition of the client lipase and its chaperone-dependent folding, but may also favour release of the folded client followed by a next chaperoning cycle, with a non-trivial effect on the resulting enzymatic activity.

To test the chaperone:client interactions as a first step, the lipase was fluorescently labelled with CF647-maleimide, and the equilibrium assembly of the LipH:LipA complex with different foldase variants was monitored through the spectral shift in the dye fluorescence emission ^28^. The interaction was resolved for all tested constructs (Figure 6A), though the affinities varied broadly. LipH^VD^ manifested high affinity to LipA, with *k*_D_ measured in low-nM range, in agreement with previous studies on the soluble chaperone ^18^, so neither the fluorescent labelling of the client lipase, nor the presence of the VD polypeptide abolished the assembly. Thus, we concluded that the VD polypeptide only transiently interacts with the chaperoning pocket, or it is displaced upon binding of the client lipase. In a striking contrast, LipH^FL^ reconstituted in either amphipols or the nanodiscs showed the affinity in the range of 200-500 nM, thus being two-orders of magnitude lower than that of the soluble LipH^VD^ (Figure 6A). Thus, the proximate environment strongly influenced the chaperone:client interactions, which has not been previously recognized.

**Figure 6.**
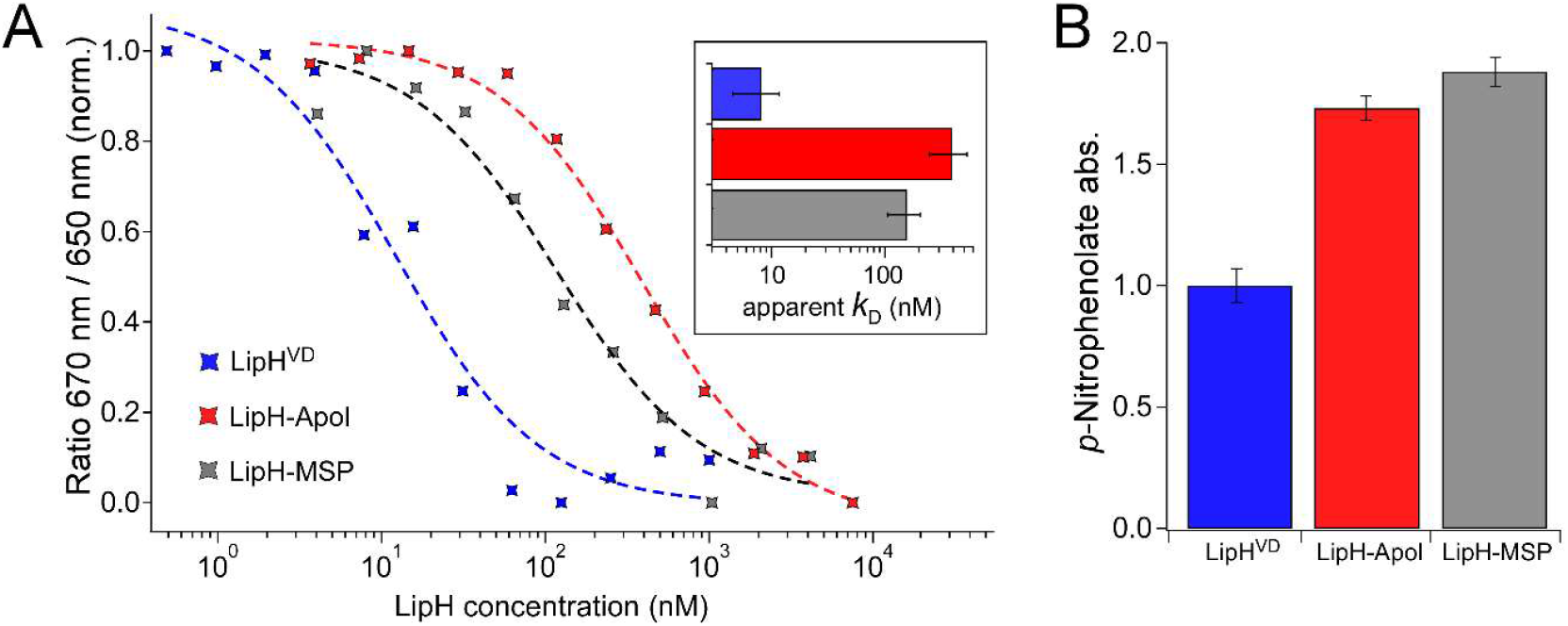
LipH:LipA assembly and activity *in vitro*. **(A)** Exemplary recordings of the spectral shift in LipA-CF647 fluorescence upon titrating LipH variants. Normalized ratios of LipA-CF647 fluorescence intensity at 670 nm and 650 nm are plotted against the LipH concentration. The apparent dissociation constants *K*_D_ calculated for each interaction as an inflection point of the fitting function (dashed lines) are shown in the inset. The error bars correspond to the standard deviations, calculated from technical triplicates. **(B)** Lipase activity *in vitro* mediated by LipH variants was determined with *p*-nitrophenylbutyrate as the substrate.

The differences in the affinity suggested that the environment of LipH^FL^ either suppressed the lipase recognition/binding, or promoted the dissociation of the complex and release of the lipase, potentially in a folded and enzymatically active state. To evaluate those scenarios, we set out to assess the LipH-dependent activity of LipA. As previously established in LipH:LipA *in vitro* studies ^5,7-9,16^, once the lipase has acquired its functional conformation, hydrolysis of a model substrate *p*NPB (para-nitrophenyl butyrate) to *p-* nitrophenolate can be monitored via a colorimetric reaction (Suppl. Figure 12), and the enzymatic activity does not require LipA release from the chaperone. In contrast, the lipase remains inactive/misfolded in absence of LipH, so only low signal associated with the substrate autohydrolysis can be observed. Hence, we implemented the enzymatic assay to compare the chaperoning properties of the soluble and the full-length LipH variants. Strikingly, both LipH-Apol and LipH-MSP constructs facilitated high activity of LipA, approx. two-fold higher than that of LipH^VD^, despite the reduced affinity (Figure 6A). Such observation strongly opposes the hypothesis that the membrane mimetics hinder the client recognition, which would result in a declined folding and activity of LipA. Instead, another scenario seems feasible: The low affinity measured for LipH^FL^:LipA originated from enhanced dissociation of the complex, so the native-like LipH^FL^ variants were able to release their folded clients into solution and to facilitate then new chaperoning rounds. Thus, a single membrane-anchored foldase could assist in folding of multiple LipA molecules, resembling the pathway taking place at the cellular membrane, while the membrane-less LipH^VD^ remained tightly bound to their clients.

### LipH MD2 region mediates tight interactions with the client lipase

Release of the lipase towards its further secretion requires rupturing interactions within the LipH:LipA complex, hence learning the dynamics of the complex would be instrumental to resolve the dissociation mechanism. The available crystal structures of the homologous complexes ^12,13^, as well as the model of LipH:LipA from *P. aeruginosa* show nearly identical tight packing of the chaperone around the folded client (Suppl. Figure 1A). Interactions of LipH MD1 are centred around the C-terminal helix of the lipase, while the MD2 contacts at least three helices of the N-terminal part of LipA (Figure 1B). To assess the potential protein dynamics within the complex, we implemented hydrogen/deuterium exchange mass spectrometry (HDX-MS) of LipH:LipA versus the individual LipH and LipA, while using the minimal functional foldase domain LipH^Chap^. HDX-MS is a valuable tool to study conformational changes, incl. folding, and protein:protein interactions, as it allows to discriminate between the solvent-exposed and buried areas of a target protein based on the polypeptide deuteration levels ^29^. By comparing HDX levels within LipA, we expected to trace its chaperone-mediated folding. In its turn, analysis focused on LipH^Chap^ could reveal how binding of the client changes the solvent exposure of the chaperone, and which areas become shielded upon this interaction. For the free LipH^Chap^ one would expect higher exposure to the solvent and rapid HDX rates, while LipA binding should suppress the exchange at the involved interfaces. With that information we may elucidate what contacts within the complex remain stable over time, and which areas may be transiently exposed due to structural fluctuations.

HDX-MS experiments showed decline in accessibility of LipA regions in presence of LipH^Chap^, (Suppl. Figure 13), in excellent agreement with the structural model of LipH:LipA complex, as the effect was most pronounced for the LipA areas covered by the chaperone. In particular the N-terminal fragment of LipA (residues 28-125, Suppl. Figure 13C and D) exhibited rapid HDX indicative for the lack of secondary structure, which was decelerated in presence of LipH^Chap^. Folding-associated HDX reduction within the C-terminus of LipA was also apparent (e.g., residues 269-311, Suppl. Figure 13C) albeit less pronounced. Thus, LipA indeed acquired its folded state and remained bound to LipH^Chap^, while in absence of the chaperone it probably formed diverse aggregates ^27 16^. For the foldase, HDX-MS analysis allowed for nearly complete coverage of the free LipH^Chap^ and the *in vitro* assembled complex LipH^Chap^:LipA (Suppl. Figure 14). Expectedly, binding of the client lipase suppressed the HDX within the chaperone, but the effect was not uniformly distributed over the structure (Figure 7A, Suppl. Figure 14C): The reduction was most pronounced within the EHD and MD2 sub-domains suggesting stable chaperone:client interactions, such as putative salt bridges between conserved residues Arg-280 and Arg-327 of LipH, and Glu-58 and Glu-66 of LipA (Figure 7B). In contrast, minor changes in the HDX levels were observed within MD1 despite the latter forming close contacts with LipA within the known structures (Suppl. Figure 1). Transient detachment of MD1 from the bound lipase or changing its position/orientation at the lipase surface would explain the high HDX rates within this region, primarily stabilized via LipH Ser-112 and LipA Gln-275.

**Figure 7.**
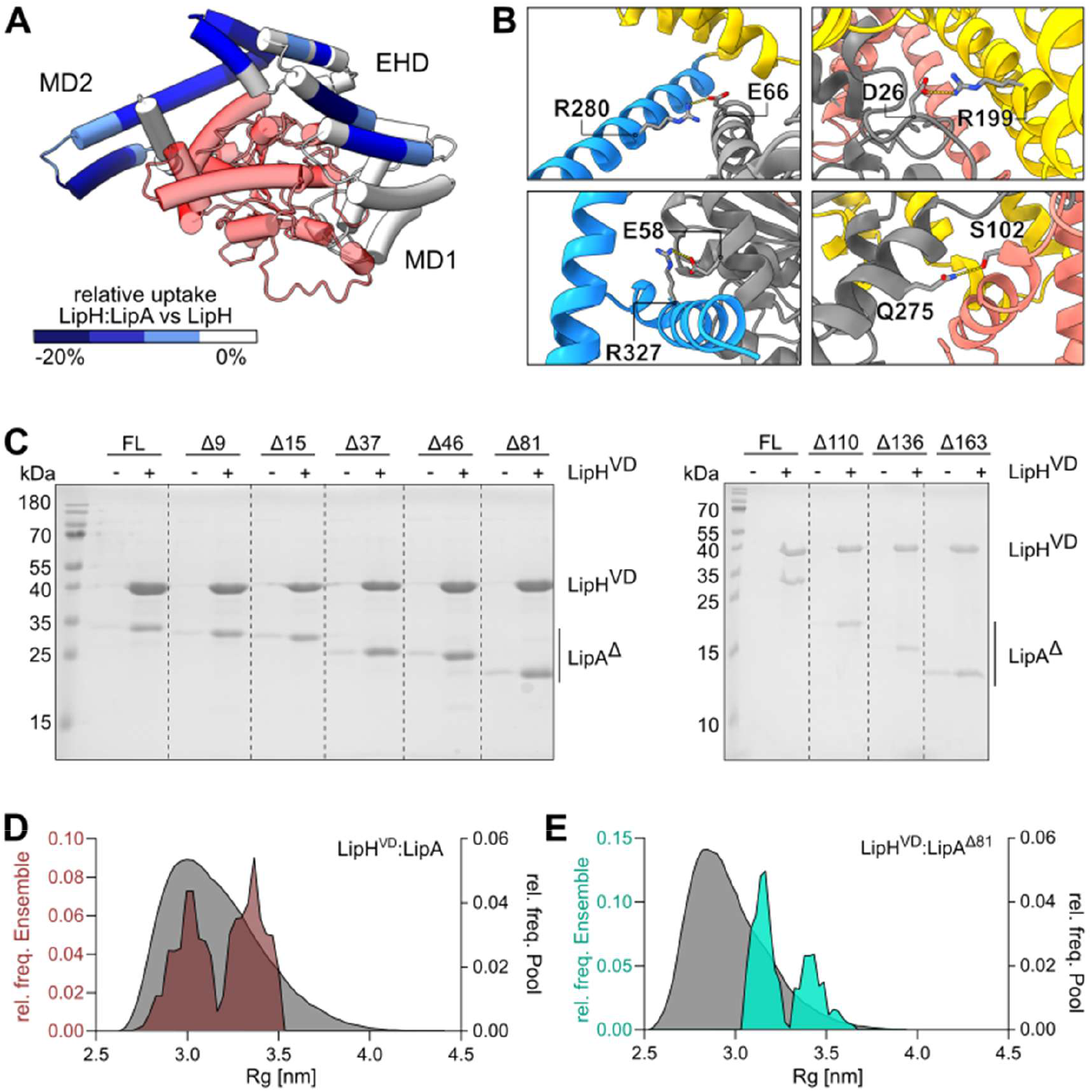
Mechanism of LipH:LipA recognition. **(A)** Changes in the hydrogen/deuterium exchange within LipH upon LipA binding plotted on the modelled LipH:LipA complex. Blue regions of LipH indicate lower deuterium incorporation of LipH in presence of LipA, and so shielding of the protein. LipA is shown in transparent red. **(B)** Electrostatic interactions between LipH and LipA predicted based on the AlphaFold3-derived model. The coloring of the LipH domains corresponds to Figure 1B; LipA is shown in grey. **(C)** Co-elution assay of LipH^VD^ and either the full-length LipA (“FL”), or the lipase fragments (“LipA^Δ^”). The number of residues deleted from the C-terminal end is indicated above each LipA fragment. For each LipA fragment, elution without and with LipH^VD^ is shown. **(D and E)** Gyration radii distribution (*R*_*g*_) from EOM of LipH^VD^:LipA and LipH^VD^:LipA^Δ81^ complexes as determined by SAXS. The distribution of the ensemble pool used for fitting is shown in grey; selected EOM models are shown in brown (LipH^VD^:LipA) and cyan (LipH^VD^:LipA^Δ81^). Wider dimensions of LipH^VD^:LipA^Δ81^ are attributed to the flexibility of the MD1 domain. Bi-modal distributions likely originate from the flexibility of the VD polypeptide.

### The N-terminal fragment of LipA carries the recognition features for chaperoning

The available structures of LipH:LipA homologs and the AlphaFold-based model of the studied complex from *P. aeruginosa* suggest multiple interactions between LipH EHD and MD2 with the N-terminal domain of the lipase (Figure 7B) ^30^. Thus, we questioned whether the chaperone:client interaction can be established with a fragment of LipA truncated at the C-terminal side, which would mimic a partially translocated lipase. To this end, we designed and isolated a series of N-terminal fragments of LipA and examined whether those are competent to interact with the polyhistidine-tagged LipH^VD^ bound to IMAC resin. Strikingly, all tested fragments co-eluted with the chaperone, including the shortest fragment LipA^Δ163^ consisting of only 122 amino acids (Figure 7C). As the elution yield dropped by approx. 80 % in absence of LipH^VD^, we concluded that LipA fragments were indeed recognized and caught by the chaperone. Notably, the urea-denatured outer membrane protein OmpA was not co-eluted with LipH^VD^ (Suppl. Figure 15), in agreement with the specificity of the LipH:client interaction.

To validate that the interaction between LipH and LipA does not essentially require the full-length lipase, we analysed the assembly of the chaperone with the lipase fragment LipA^Δ81^, as well as the full-length LipA, using SEC-SAXS (Suppl. Figures 16 and 17). The measured molecular masses of LipH^VD^:LipA^Δ81^ (58.2 kDa) and LipH^VD^:LipA (67.1 kDa) closely matched the calculated values of 59.6 kDa and 68.5 kDa, respectively. Thus, the chaperone:client complexes were indeed formed in 1:1 stoichiometry, and the N-terminal fragment of LipA was sufficient for binding (Suppl. Table 1). For both complexes, Kratky plots showed bell-shaped profiles indicating compact and folded molecules (Suppl. Figures 16C and 17C), which contrasted the broader profile seen for LipH^VD^ alone (Suppl. Figure 10C). Thus, the flexibility of the chaperone substantially decreased upon binding the client. Interestingly, even though the LipH^VD^:LipA^Δ81^ complex had a lower molecular weight, it manifested a larger radius of gyration than that of LipH^VD^:LipA, 3.53 nm vs. 3.26 nm, as provided by the pair-distance distribution function *p(r)* (Suppl. Table 1). Thus, LipH^VD^:LipA^Δ81^ acquired more extended conformation than LipH^VD^:LipA, which could be explained by missing interactions of LipH MD1 with the C-terminal part of LipA. In agreement, fitting the SAXS data with structural models of the chaperone:client complex via the EOM approach resulted in bimodal Rg distribution behaviour for both complexes (Figure 7D and E; Suppl. Table 1), but a more relaxed distribution was observed for LipH^VD^:LipA^Δ81^ in comparison to LipH^VD^:LipA. Taken together, the results suggest that the N-terminal fragment of the lipase carries the recognition features that stabilise the complex in a more compact conformation.

## Discussion

Chaperone-assisted protein folding has been a topic of extensive research over decades, which has offered detailed insights on the universally conserved families of ATP-dependent chaperones, such as Hsp70/40, or energy-independent bacterial chaperones, such as trigger factor, SurA, and Skp ^31,32^. Those chaperones constitute essential parts of cellular proteostasis, as they participate in a global control of the protein content via balancing protein folding and degradation. To ensure their broad specificity, the recognition of clients is based on basic physico-chemical principles, such as exposure of hydrophobic polypeptide stretches or aromatic residues, that is characteristic for incomplete or aberrant folding states ^33^. Lack of specific fine-tuned interactions results in weak affinities, being typically in the range of tens of µM, so one client protein may subsequently interact with several chaperones prior being either correctly folded or routed for degradation ^34^.

Differently to those, several classes of chaperones with narrow specificity have been described in bacteria, with examples of type III secretion system adaptor components ^35^, and steric foldases. While the T3SS adaptors are widespread and diverse, only two classes of steric foldases have been identified so far, which facilitate folding of either bacterial lipases or type II proteases, both targeted then for secretion via T2SS ^36,37^. The known steric foldases share a primary topology, as their periplasm-exposed chaperoning domains are anchored via transmembrane helices and unstructured linker polypeptides to the lipid bilayer, so the chaperones are not co-secreted together with their clients. On example of *P. aeruginosa* LipH, our MD simulations and SAXS analysis show here that the unstructured linker allows extensive motions of the chaperoning domain, so the membrane-anchored LipH is able to sample a large area within the periplasm to bind the client lipase, though the linker may transiently occlude the binding interface. Notably, the foldases with the unstructured linkers clearly differ from the well-studied membrane-anchored chaperone complex PpiD-YfgM that contains short and structured domains (Suppl. Figure 18) ^38,39^. The PpiD-YfgM assembly interacts with the SecYEG translocon, so the rigid linkers may optimally position the chaperoning moiety for screening and/or capturing the newly translocated clients.

Remaining in the proximity to the membrane, the chaperone domain of LipH can interact with the lipid leaflet, and the simulations suggest that the chaperone accessibility depends on the environment: An elevated ionic strength allows interactions of the chaperoning domain with the membrane interface, which may negatively affect the client capture. Interestingly, our previous work showed that at such conditions the chaperone Skp efficiently rescues LipA from aggregation in solution and supports its secretion *in vivo*, so Skp can serve as a holdase chaperone for the lipase and provide additional timing for LipH to bind its specific client, thus minimizing the off-route misfolding pathways ^16^.

Both MD simulations of LipH^FL^ and SAXS analysis of the soluble LipH^VD^ confirm flexibility within the chaperoning domain, where the mini-domains MD1 and MD2 move at the opposing sides of the bridging EHD. These findings qualitatively corroborate the results of an earlier study, where simulations and single-molecule FRET experiments revealed a continuum of conformations explored by LipH^Chap 18^. Those movements, together with the dynamics of the VD polypeptide, result in a wide ensemble of LipH configurations which differ from the defined conformation seen in the foldase:lipase structure, raising a question about the recognition mechanism. Our biochemical and structural analyses suggest for the first time that the recognition may not involve the complete LipA, but shorter fragments may be captured, first of all at the MD2-EHD interface of LipH, followed by loading of a complete lipase into the chaperoning cavity for folding and activation. Hypothetically, such recognition could occur already during LipA translocation via the SecA:SecYEG machinery, once the N-terminal tail emerges at the periplasmic side, followed by gradual loading coupled with the elongation of the LipA polypeptide chain.

While LipH remains flexible upon recognition of the LipA fragment, SAXS analysis shows that the LipH dynamics is suppressed once the chaperone binds the full-length LipA. Notably, the HDX-MS data reveal low to no shielding of the MD1 domain by the lipase, implying that the interactions formed by MD1 are marginally stable. Indeed, a single mutation within this region drastically affects the MD1:LipA interactions, but also the chaperoning properties of LipH ^40,41^. Thus, it is tempting to speculate that the LipH MD2 domain plays a leading role in recognition and binding of the client, while MD1 contributes to the chaperoning activity, such as correct positioning of LipA helices, and, possibly, release of the correctly folded lipase.

The mechanism of the client release remains another open question in our understanding of the steric chaperones. Our data, as well as previous studies, demonstrate high, nanomolar affinity of the LipH chaperoning domain to the lipase, which may not be compatible with spontaneous release of LipA and its handover to Xcp-T2SS. Release of clients from the non-specific periplasmic chaperone Spy has been recently linked to its disordered N-terminal end: Electrostatic interactions of the polypeptide with the binding pocket of Spy serve to expel client proteins ^42^. In contrast, the non-conserved VD polypeptide of LipH^VD^ did not affect the affinity for the lipase, despite its transient interactions with the chaperoning domain. This finding speaks against its direct role in destabilization of the complex. However, our functional analysis of the full-length chaperone embedded in nanodiscs and amphipols suggested that the membrane mimetics may stimulate release of the client lipase: While LipH retained its chaperoning properties, the affinity to LipA dropped manifold upon reconstitution. We hypothesize that the anchored LipH^FL^ facilitated multiple chaperoning rounds associated with binding and release of the client lipase, thus resembling the pathway occurring in the cell. One potential stimulus for LipA release could be the electrostatic properties of the proximate environment. Both the phospholipid-based nanodiscs and the carboxyl-rich amphipol A8-35 are the highly polar and negatively charged “carriers”, which may either modulate dynamics of the LipH MD1 domain or may interact directly with the folded lipase, which exposes its cationic side to the solvent (Suppl. Figure 19).

To summarize, our study offers the first comprehensive view on the conformational dynamics of the lipase-specific foldase LipH at the membrane interface, and suggests that the foldase:lipase recognition is mediated by specific regions of both proteins, i.e. MD2 of LipH and the N-terminal fragment of LipA, while MD1 dynamics serves for the lipase folding and release (Figure 8). It remains to be tested though whether the molecular mechanism derived from *in vitro* and *in silico* analysis can be recapitulated in the living cells. Our results provide a solid ground for such studies, which may test potential effects of specific point mutations at the LipH:LipA interface on the lipase secretion *in vivo*. Once successful, the chaperone-dependent folding of the lipase may be modulated at various stages to suppress secretion of a virulence factor in *P. aeruginosa* and related species, but also to facilitate production of the functional lipase in biotechnological applications.

**Figure 8.**
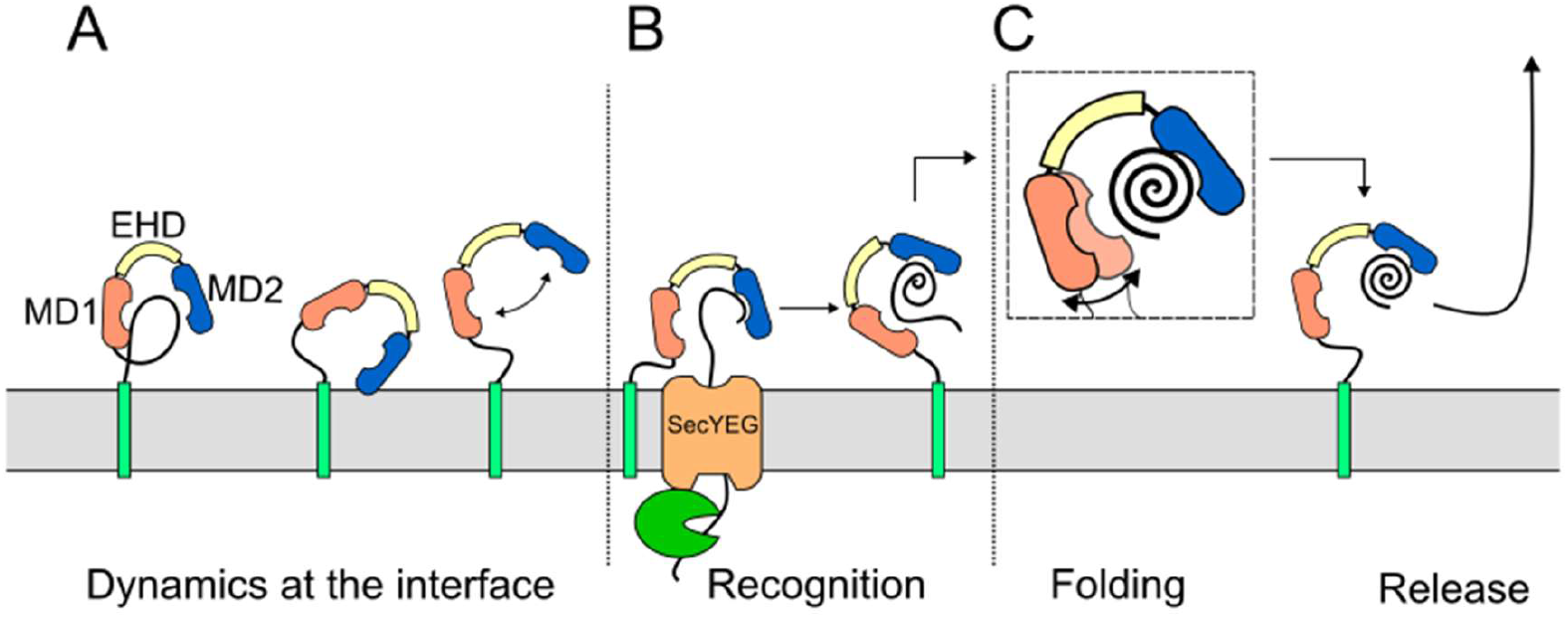
LipH conformational dynamics and interactions with the client lipase. **(A)** The foldase LipH exhibits distinct dynamic behaviours at the membrane interface, ranging from transient contacts of the unstructured linker polypeptide to breathing motions of the chaperoning domain, including opening and closing transitions, and membrane interactions. **(B)** Recognition and binding of the N-terminal fragment of the client lipase is facilitated by the MD2 domain of LipH (blue), potentially at the translocation stage. **(C)** MD1 (salmon) contributes predominantly to the chaperoning process, including proper positioning of LipA helices, and weak interactions in this region potentially facilitate the release of the folded lipase.

## Acknowledgements

We would like to thank Lucas Kühl and Dr. Florian Altegoer for the help with the analysis of LipH homologs and bioinformatics, Dr. Cristian Rosales-Hernandez for the mass photometry measurements, and Dr. Filip Kovacic for many discussions on the topic. We acknowledge the computational infrastructure and support provided by the Centre for Information and Media Technology (HHU Düsseldorf), and Prof. Dr. Birgit Strodel to providing the computing time on the supercomputer JURECA (Forschungszentrum Jülich). We acknowledge DESY (Hamburg, Germany), a member of the Helmholtz Association HGF, for the provision of experimental facilities. Parts of this research were carried out at PETRA III, and we would like to thank Tobias Gräwert and Cy M. Jeffries (EMBL Hamburg) for their assistance in using the beamline P12. We acknowledge the European Synchrotron Radiation Facility (Grenoble, France) for provision of synchrotron radiation facilities, and we would like to thank Mark Tully for his assistance in using beamline BM29.

The study was funded by the German Research Foundation (Deutsche Forschungsgemeinschaft, DFG) within the Collaborative Research Center 1208 “Identity and Dynamics of Membrane Systems” (grant number 267205415 to A.K. and K.E.J.) and supported through the Marburg core facility for Interaction, Dynamics and Assembly of Biomolecular Structures (DFG grant numbers 324652314 and 260989694 to Prof. Gert Bange, Marburg). The Center for Structural Studies is part of StrukturaLINK Rhein-Ruhr which is funded by the DFG (grant number 573727698 and 417919780) and INST 208/761-1 FUGG.

## Data availability

The SAXS data have been deposited to the Small Angle Scattering Biological Data Bank ^43^, with the accession codes SASDYM9 (LipH^VD^), SASDYN9 (LipH^VD^:LipA) and SASDYP9 (LipH^VD^:LipA^Δ81^). HDX-MS data have been deposited to the ProteomeXchange Consortium via the PRIDE partner repository ^44^ with the dataset identifier PXD074746.

## Supporting information

This article contains supporting information.

## Methods

### Bioinformatics analysis of the LipH variable domain

To identify the LipH homologs across bacteria families, the EFI - Enzyme Similarity Tool with default options ^45^ was used to find sequences similar to the one of *Pseudomonas aeruginosa* LipH (pseudomonas.com: PA2863). The obtained protein accession numbers were added to Uniprot ID mapping server to get the corresponding FASTA data, which were further analysed via InterPro ^46^ using the tools from Superfamily ^47^ and TMHMM ^48^. Proteins containing both the N-terminal hydrophobic anchor and the Lif-like domains were used for further analysis, with the final count of 858 sequences. The polypeptide chain between the predicted anchor and the Lif-like domain corresponded to the “variable domain” region of LipH, and the length and the amino acid composition was analysed.

### Expression and purification of LipH^VD^

The gene encoding for the soluble LipH variants, which lack the residues 5 to 22 (LipH^VD^) and 5 to 62 (LipH^Chap^) of the full-length foldase, were cloned into pET21a vector as previously described ^16^. *E. coli* BL21(DE3) cells transformed with the respective plasmid were grown in LB medium supplemented with 100 μg/mL ampicillin at 37°C while shaking at 180 rpm. Upon reaching OD_600_ of 0.6, LipH expression was induced by adding 0.5 mM isopropyl-β-D-thiogalactopyranoside (IPTG; Merck/Sigma-Aldrich) and carried out for 2.5 h. Cells were harvested and lysed using a microfluidizer (M-110P, Microfluidics Corp.), and the debris and membranes were removed by ultracentrifugation at 42000 rpm for 1 h (45Ti rotor, Beckman Coulter). For further purification via immobilized metal ion affinity chromatography (IMAC), the supernatants were incubated with Ni^2+^-NTA agarose resin (Protino, Macherey-Nagel GmbH & Co. KG) for 1 h at 4°C. The resin was washed with 50 mM Hepes-KOH pH 7.4, 500 mM KOAc, 5 % glycerol (v/v), 10 mM imidazole, and the target protein was eluted with 50 mM Hepes-KOH pH 7.4, 150 mM KOAc, 5 % glycerol, 200 μM, 300 mM imidazole. The elution fractions were concentrated and subjected to size exclusion chromatography (SEC) in 50 mM Hepes-KOH pH 7.4, 150 mM KOAc, 5 % glycerol, using Superdex 200 Increase GL 10/300 column with ÄKTA pure set-up (Cytiva). The protein concentration was determined spectrophotometrically (NeoDot, NeoBiotech) using calculated extinction coefficient of 19 940 M^-1^ cm^-1^ for both variants.

### Expression and purification of the full-length LipH

The gene encoding for the full-length foldase LipH^FL^ was cloned into pBAD vector via KpnI and HindIII restriction sites to contain an N-terminal octa-histidine tag. *E. coli* BL21(DE3) cells transformed with the respective plasmid were grown in LB medium supplemented with 100 μg/mL ampicillin at 37°C while shaking at 180 rpm. Upon reaching OD_600_ of 0.6, LipH^FL^ expression was induced by adding 0.2% arabinose and carried out for 2 h. The cells were harvested by centrifugation at 5000 *g* for 20 min at 4°C (rotor SLC-6000, Sorvall/Thermo Fisher) and subsequently resuspended in 20 mM Tris (hydroxymethyl)aminoethane-HCl pH 8.0, 100 mM NaCl, 1 mM DTT (Merck/Sigma-Aldrich), 5 % glycerol and cOmplete protease inhibitor cocktail (Roche). Cells were lysed using microfluidizer (M-110P, Microfluidics Inc.), and the cell debris was removed by centrifugation at 18000 *g* for 10 min at 4°C (rotor SS34, Sorvall/Thermo Fisher). The supernatant was then centrifuged at 205000 *g* for 1 h at 4°C (rotor 45 Ti, Beckman Coulter) to pellet crude membranes. The membranes were resuspended in the same buffer (20 mM Tris-HCl pH 8.0, 100 mM NaCl, 1 mM DTT, 5 % glycerol, and cOmplete protease inhibitor cocktail) and stored at - 80°C until further usage. To extract LipH^FL^, n-dodecyl-β-D-maltopyranoside (DDM; Glycon Biochemicals GmbH), n-decyl-β-maltoside (DM; Glycon Biochemicals GmbH), lauryl maltose neopentyl glycol (LMNG; Anatrace) and 6-cyclohexyl-1-hexyl-β-D-maltoside (Cymal-6; Anatrace) were used. Solubilization was performed with 1 % of each detergent in 50 mM Tris-HCl pH 8.0, 100 mM NaCl, 0.2 mM TCEP, 5 % glycerol, and cOmplete protease inhibitor cocktail for 1 h at 6°C on a rolling table. After solubilization, the samples were centrifuged at 20.000 *g* for 10 min at 4°C (Hermle Z216 M) to remove non-soluble material. The his-tagged LipH^FL^ were immobilized on Ni^2+^-NTA agarose resin equilibrated with the wash buffer (50 mM Tris-HCl pH 8.0, 100 mM NaCl, 0.2 mM TCEP, 20 mM imidazole, cOmplete protease inhibitor cocktail) supplemented with the detergent of choice. The detergent concentrations for the washing and elution steps were ∼5-fold above the specific critical micelle concentrations (CMC) resulting in 0.05 % (w/v) for DDM, 0.4% for DM, 0.005% for LMNG and 0.1% for Cymal-6. The resin was washed extensively with the wash buffer to remove weakly bound impurities, and the protein was eluted with 50 mM Tris-HCl pH 8.0, 100 mM NaCl, 0.2 mM TCEP, 300 mM imidazole, 10 % glycerol, cOmplete protease inhibitor cocktail. For size-exclusion chromatography, the protein was loaded on Superdex 200 Increase 10/300 GL column (Cytiva) in the desired buffer compositions (e.g., 50 mM Tris-HCl pH 8.0, 100 mM NaCl, 0.2 mM TCEP, 10 % glycerol, 0.005 % LMNG). The concentration of the purified LipH^FL^ was determined spectrophotometrically and the purification yield was further analysed via SDS-PAGE.

### LipH^FL^ reconstitution into membrane mimetics

For the amphipol reconstitution, 15 nmol LipH^FL^ in DDM was mixed with three-fold excess of amphipol A8-35 (Anatrace) and incubated for 2 h while rolling at 6 °C. To remove the detergent, 60 mg washed and dried Bio-Beads SM-2 sorbent (Bio-Rad Laboratories GmbH) was added to the mixture and incubated overnight while rolling at 6° C. The reconstitution reaction was loaded on Superdex 200 Increase 10/300 GL, and the fractions were analysed via SDS-PAGE. For the nanodisc reconstitution, membrane scaffold protein MSP1E3D1 was purified as described before ^24^. Liposomes were prepared of 70% 1-palmitoyl-2-oleoyl-glycero-3-phosphocholine (POPC) and 30% 1-palmitoyl-2-oleoyl-sn-glycero-3-phospho-(1-rac-glycerol) (POPG) (Avanti Polar Lipids, Inc). Lipids were mixed from chloroform stocks to achieve the target POPC:POPG ratio, and the solvent was evaporated at 200 mbar and 40°C for 30 min. The film of dried lipids was resuspended in 50 mM Hepes/KOH pH 7.4, 50 mM KCl and extruded through a porous polycarbonate membrane (200 nm) using the Mini-Extruder set (Avanti Polar Lipids, Inc). Subsequently, the liposomes were dissolved using 0.5% Triton X-100. LipH^FL^ purified in DDM was reconstituted into nanodiscs at the protein:MSP:lipid molar ratio of 1:6:600. To remove the detergent, 60 mg washed and dried Bio-Beads SM-2 sorbent was added to the mixture and incubated overnight while rolling at 4°. The empty and loaded nanodiscs were separated by size exclusion chromatography using Superdex 200 10/300 Increase GL column, and the fractions were analysed via SDS-PAGE. Mass photometry analysis of the reconstituted LipH^FL^ samples was performed using Two^MP^ instrument (Refeyn Ltd.) and calibrated using NativeMark Unstained Protein Standard (Invitrogen/Thermo Fisher Scientific).

### Expression and purification of LipA

The mature lipase LipA^F144E^ (pseudomonas.com:PA2862) lacking the signal peptide (residues 1 to 27), and its C-terminal truncations were expressed and isolated as previously described ^16^. The mutation F144E within the lid domain reduces the aggregation propensity of the lipase ^16^. For the C-terminal truncations, the amino acids 300-309 (Δ9), 294-309 (Δ15), 272-309 (Δ37), 263-309 (Δ46), 228-309 (Δ81),199-309 (Δ110), 173-309 (Δ136) and 146-309 (Δ163) were removed via PCR with subsequent blunt end ligation. *E. coli* BL21(DE3) cells were transformed with a corresponding pET22b-based plasmid and grown at 37°C shaking at 180 rpm. Overexpression was induced by addition of 0.5 mM IPTG and conducted for 2 h. Afterwards, the cells were harvested by centrifugation at 5000 *g* at 4°C for 20 min. The cell pellet was resuspended in *E. coli* Lysis reagent (New England Biolabs GmbH) and incubated at 20°C for 15 min. The inclusion bodies were pelleted upon centrifugation and repeatedly washed with 20 mM Tris-HCl pH 7.5. After three rounds of washing, the inclusion bodies were dissolved in 8 M urea, 50 mM Tris-HCl pH 7.5. LipA constituted above 90% of the inclusion bodies content, as determined via SDS-PAGE and following staining (Quick Coomassie stain, Serva).

### In vitro *activity of LipA*

The hydrolytic activity of LipA was measured *in vitro* using the model substrate *para*-nitrophenyl butyrate (*p*NPB) as described elsewhere ^16^. For the measurement, 2 μM of the urea-denatured LipA was mixed with an equimolar amount of the soluble chaperone variant LipH^VD^ or the full-length foldase LipH^FL^ embedded into nanodiscs or amphipols. Due to the intrinsic absorbance of MSP1E3D1, the concentration of LipH^FL^ in nanodiscs was estimated colorimetrically based on the band intensity on SDS-PAGE (Amersham 600 RGB imager, ImageQuant software, both Cytiva). The total volume of the reaction was 40 μL, adjusted with TGCG buffer (5 mM Tris-HCl pH 9.0, 5 mM glycine, 1 mM CaCl_2_, and 5 % glycerol). The samples were incubated for 15 min at 37°C to allow for LipH:LipA complex formation. Afterwards, 10 μL sample were transferred into a 96-well plate containing 100 μL TGCG. *p*NPB stock was diluted to 10 mM in acetonitrile, and it was further diluted 10-fold with 50 mM triethanolamine pH 7.4 prior the measurement. 100 μL of the *p*NPB solution was pipetted to each well for measurement and the reaction was carried out at 37°C up to 3 h, while monitoring the absorbance of the hydrolysis product, *p*-nitrophenolate, at 410 nm (Infinite 200 PRO plate reader, Tecan). Absorbance values within the exponential phase (900 sec) were used to compare the hydrolysis levels at different conditions. Loading controls were performed for all checked conditions and the intensity of bands on the SDS-PAGE was quantified (ImageQuant, Cytiva). The obtained values were corrected for variations in LipA as well as the chaperone after deduction of autohydrolysis. The measurements were conducted in biological triplicates, each in technical quintuplicates.

### Spectral shift measurements

The interactions of LipH variants with LipA were probed via the fluorescence spectral shift technique. LipA was labelled with the far-red dye CF647-maleimide (Biotium Inc.) via coupling to the endogenous cysteines. Prior to each measurement, the urea-denatured LipA-CF647 was diluted with buffer containing 5 mM Tris-HCl pH 9.0, 5 mM glycine, 1 mM CaCl_2_, 5 % glycerol, 0.5 mg/mL BSA,0.05% (w/v) Tween-20. LipA-CF647 concentration was set to 20 nM in experiments with LipH^VD^ and LipH-MSP, and 10 nM for LipH-Apol. LipH concentrations ranged from 0.5 nM to 7.5 µM. Fluorinated octyl maltoside (Anatrace) was added at concentration of 0.05% to the reaction with LipH-MSP to prevent the sample aggregation at the capillary surface (Monolith premium Capillaries, MO-K025, Nanotemper Technologies GmbH). The experiments were performed at Monolith.X instrument (Nanotemper Technologies GmbH). The excitation power was set to 40%, with medium infra-red laser power, and the fluorescence spectral shift of the LipA-bound dye between wavelengths of 670 nm and 650 nm was measured. The LipH-dependent shift values were fitted using a Hill equation, with a Hill coefficient of 1, and providing an estimate for the LipH:LipA dissociation constant.

### Co-elution interaction assay

To test interactions between the soluble foldase variant LipH^VD^ and LipA fragments, an IMAC-based co-elution assay was designed. 50 mL Ni^2+^-NTA agarose resin was loaded into low-binding reaction tubes (Sarstedt), and 30 μM of the hexa-histidine-tagged LipH^VD^ was added to the resin. Only buffer was added for reference reactions. The samples were mixed by vortexing on Vortex genie 2 (NeoLab) for 5 sec. The reaction tubes were placed on a rolling bench for 5 minutes at 4°C to let the chaperones bind to the resin. After this, 400 μL of buffer consisting of 5 mM Tris, 5 mM glycine, 1 mM CaCl_2_ was added, supplemented with 5 mM imidazole to reduce unspecific binding (wash buffer). Then an equimolar amount of LipA was added and left for 30 minutes on the rolling bench at 4°C. Afterwards the resin was pelleted for 30 s at 20.0000 g in tabletop centrifuge (Hermle Z 216 M) and the supernatant was discarded. After two rounds of washing/pelleting steps, 100 μL elution buffer containing 300 mM imidazole was added, the tubes were vortexed and incubated for 10 min on the rolling bench at 4°C. The resin was pelleted, and the supernatant with the eluted material was carefully collected. 10 µL of the elution samples were loaded on SDS-PAGE.

### Small-angle X-ray scattering

SAXS data from LipH^VD^ were collected on a Xeuss 2.0 Q-Xoom system (Xenocs), equipped with a PILATUS 3 R 300K detector (Dectris) and a GENIX 3D CU Ultra Low Divergence X-ray beam delivery system. The chosen sample-to-detector distance for the experiment was 0.55 m, resulting in an achievable q-range of 0.05-5.5 nm^-1^. The measurement was performed at 10°C with a protein concentration range up to 11.8 mg/mL. The samples were injected into the Low Noise Flow Cell (Xenocs) via autosampler, and 24 frames were collected with an exposure time of 10 min/frame. SAXS data from the LipH^VD^:LipA complex were collected on the P12 beamline (PETRA III, DESY Hamburg ^49^). The sample-to-detector distance of the P12 beamline was 3.00 m, resulting in an achievable q-range of 0.03-7.0 nm^-1^. For complex formation of LipH^VD^ and LipA, both proteins were mixed in the given ratio and preincubated for 30 min at 37°C. Prior to injection, the samples were centrifuged for 15 min at 20.000 x g. SEC-SAXS of the LipH^VD^:LipA complex was performed at 20°C on a Superdex 200 Increase 10/300 GL column (100 µL inject, buffer: 5 mM Tris pH 8.0, 5 mM glycine,1 mM CaCl_2_, 5% glycerol) with a flow-rate of 0.6 mL/min. 2400 frames were collected with an exposure time of 0.995 sec/frame. SEC-SAXS data from LipH^VD^:LipA^Δ81^ complex was collected on beamline BM29 at the ESRF Grenoble ^50^. The BM29 beamline was equipped with a PILATUS 2M detector (Dectris) at a fixed distance of 2.827 m, resulting in an achievable q-range of 0.025-5.5 nm^-1^. SEC-SAXS was performed at 20°C on a Superdex 200 increase 10/300 GL column (100 µL inject, buffer: 5 mM Tris pH 8.0, 5 mM glycine,1 mM CaCl_2_, 5% glycerol) with a flow-rate of 0.6 mL/min. 1200 frames were collected with an exposure time of 2 sec/frame. All collected data were scaled to absolute intensity against water.

Programs used for data processing were part of the ATSAS Software package ver. 3.0.5 ^51^. Primary data reduction was performed with the programs CHROMIXS ^52^ (for SEC-SAXS) and PRIMUS ^53^.With the Guinier approximation ^54^, we determine the forward scattering *I(0)* and the radius of gyration (*R*_*g*_). The program GNOM ^55^ was used to estimate the maximum particle dimension (*Dmax*) with the pair-distribution function p(r). Structural models of LipH in *apo-*state, as well as in complex with LipA variants were built via AlphaFold3 ^56^. For a better agreement with the experimental scattering data in *apo*-state, EOM was used to model the N-terminal polypeptide of LipH and to allow movement of the MD1, EHD and MD2 domains ^57,58^. For the complexes formed with LipA and LipA^Δ81^, only flexibility of LipH N-terminal end was allowed.

### Hydrogen/deuterium exchange mass spectrometry

Protein sample stocks for HDX-MS contained 50 µM of either individual LipA or LipH^Chap^, or both proteins premixed to form the LipH:LipA complex, and preparation of individual samples was aided by a two-arm robotic autosampler (LEAP Technologies) essentially as previously described ^59^. In short, HDX reactions were initiated by 10-fold dilution of the protein stock solutions with TGC buffer (5 mM Tris-HCl pH 9.0, 5 mM glycine, 1 mM CaCl_2_) prepared in D_2_O and incubated for 10, 100, 1,000 or 10,000 s at 25 °C. The HDX reaction was stopped upon adding an equal volume of quench buffer (400 mM KH_2_PO_4_/H_3_PO_4_, 2 M guanidine-HCl; pH 2.2) pre-dispensed and thermostated at 1 °C. Then, 100 μL of the resulting mixture was injected into an ACQUITY UPLC M-Class System with HDX Technology ^60^. Non-deuterated samples were generated by the same procedure through 10-fold dilution with TGC buffer prepared with H_2_O. The injected HDX samples were washed out of the injection loop (50 μL) with water supplemented with 0.1% (v/v) formic acid at 100 μL/min flow rate and guided to a column containing immobilized porcine pepsin thermostated at 12 °C. The resulting peptic peptides were collected on a trap column (2 mm x 2 cm), that was filled with POROS 20 R2 material (Thermo Scientific) and kept at 0.5 °C. Digestion and trapping was conducted for 3 min, the trap column was placed in line with an ACQUITY UPLC BEH C18 column (1.7 μm, 1.0 x 100 mm; Waters), and peptides eluted with a gradient of water supplemented with 0.1% (v/v) formic acid (eluent A), and acetonitrile supplemented 0.1% (v/v) formic acid (eluent B) at the flow-rate of 30 μL/min as follows: 0-7 min/95-65% A, 7-8 min/65-15% A, 8-10 min/15% A.

Eluting peptides were guided to a Synapt G2-Si mass spectrometer (Waters) and ionized with by electrospray ionization (capillary temperature 250 °C, spray voltage 3.0 kV). Mass spectra were acquired with the software MassLynX MS version 4.1 (Waters) over a range of 50 to 2,000 *m/z* in enhanced high-definition MS (HDMS^E^) ^61,62^ or high-definition MS (HDMS) mode for non-deuterated and deuterated samples, respectively. Lock mass correction was conducted with [Glu1]-Fibrinopeptide B standard (Waters). During separation of the peptides on the C18 column, the pepsin column was washed three times by injecting 80 μL of 0.5 M guanidine hydrochloride in 4% (v/v) acetonitrile. Blank runs (injection of double-distilled water instead of protein sample) were performed between each sample. All measurements were carried out in triplicates.

Peptides were identified and evaluated for their deuterium incorporation with the softwares ProteinLynx Global SERVER 3.0.1 (PLGS) and DynamX 3.0 (both Waters) as previously described ^59^ employing the amino acid sequences of LipA, LipH^Chap^, and porcine pepsin as database. Whenever possible, multiple charge states were utilized. All spectra were manually inspected and omitted if necessary, e.g. in case of low signal-to-noise ratio or the presence of overlapping peptides disallowing the correct assignment of the isotopic clusters. Raw data of deuterium uptake by the identified peptides and residue-specific HDX are provided in Supplemental Dataset 1.

### Molecular dynamics simulations

All molecular dynamics (MD) simulations carried out in this study are summarized in Table 1. The systems were built via CHARMM-GUI ^63-65^ by using the force field CHARMM36m and TIP3P ^66^ water model, with the temperature set at 298 K (25°C), the pressure - at 1 bar, and pH 7.0. The membrane was composed of di-oleoyl-phosphatidylethanolamine (DOPE; 75 mol %), di-oleoyl-phosphatidylglycerol (DOPG; 20 mol %) and 1’,3’-bis{18:1/18:1-sn-glycero-3-phospho]-glycerol (cardiolipin, CL; 5 mol %), thus mimicking the bacterial inner membrane at two different ion conditions (150 mM and 25 mM NaCl). To perform the simulations and analysis, we used MD software package GROMACS 2021/2022^67^.

Derived on the AlphaFold-based model of the full-length LipH, the N-terminal helix was inserted in the membrane via CHARMM-GUI ^63-65^. The lipid bilayer had lateral dimensions of 22 x 22 nm^2^ (x/y-plane), and it was placed in the middle of the simulation box with the total height of 19 nm (z-axis), resulting in a system size of ∼900,000 atoms. The *mdp* scripts were used for energy minimization, equilibration in five steps and production run from CHARMM-GUI web server since they are optimized for protein-membrane simulations. The energy was minimized to 1,000 kJ*mol^-1^*nm^-1^ using the steepest descent algorithm, followed by five-step equilibration to the desired temperature of 298 K (25°C) and pressure of 1 bar. First, two *NVT* equilibration steps were applied to keep constant the number of atoms (*N*), the box volume (*V*), and temperature (*T*), followed by three-step *NpT* equilibration to adjust the pressure (*p*). The protein’s and lipid’s heavy atoms were restrained to allow the water molecules and ions to relax around the solute but they were decreased by every equilibration step. The Berendsen thermostat ^68^ was employed to regulate the temperature in the *NVT* simulations, while the Berendsen thermostat and the semi–isotropic Berendsen barostat ^68^ were employed for the *NpT* simulations. The PME method ^69,70^ was applied to calculate long-range electrostatic interactions with periodic boundary conditions. Van der Waals and Coulombic interaction cut-offs were set to 1.2 nm using the LINCS algorithm ^71^ to constrain all bond-lengths to hydrogens. MD runs were performed for 500 ns with a time step of 2 fs by recording the coordinates and velocities every 20 ps as well as the Nosé-Hoover thermostat ^72,73^ and the semi– isotropic Parrinello-Rahman barostat ^74^.

To follow possible changes in the secondary structure of LipH during the MD simulations, DSSP algorithm ( for *Define Secondary Structure of Proteins*) was employed ^75^. The contact map shows the intramolecular interactions within chaperone domain of LipH as percentage during the simulation. Here, a cut-off value under 6 Å was applied to consider as contact, which is depicted in a blue-white-red colour scheme, where red colours represent higher contact percentage. The DSSP and contact maps were calculated via MDanalysis 2.9.0 ^76^ and own python script. For DSSP analysis, the secondary structure was classified as helical (α-, 3_10_ and π), β-sheets/β-strands, and coil that summarized other elements provided by the algorithm (coil, bend, turn). To assess the dynamics of the chaperoning domain of LipH, the radius of gyration and the minimal distance between residues 107 (MD1) and 321 (MD2) was analyzed.

To visualize positions of LipH in the simulation box during MD simulations, the density maps of ions and water were assembled via *gmx densmap*. Here, the direction of the y-axis was used to integrate the signal, the binning size was set to 0.08 and the unit was molecule per nm^3^. DuIvyTools (https://github.com/CharlesHahn/DuIvyTools) were used to convert the format from *xpm* to a matrix *dat* file (*dit xpm2dat*), as required for plotting the density maps via an own python script. The minimal distance between the membrane and LipH was computed for the whole protein, as well as the different domains to characterize LipH:membrane interactions. If not stated otherwise, all analysis were computed with GROMACS 2021/2022, and data was plotted with Gnuplot. PyMol ^77^ was used for rendering structural figures related to MD simulations.

