## Supplemental Figures & Table for "Conformational dynamics of the membrane-anchored foldase LipH from *Pseudomonas aeruginosa* governs recognition and release of its client lipase"

### Supporting information

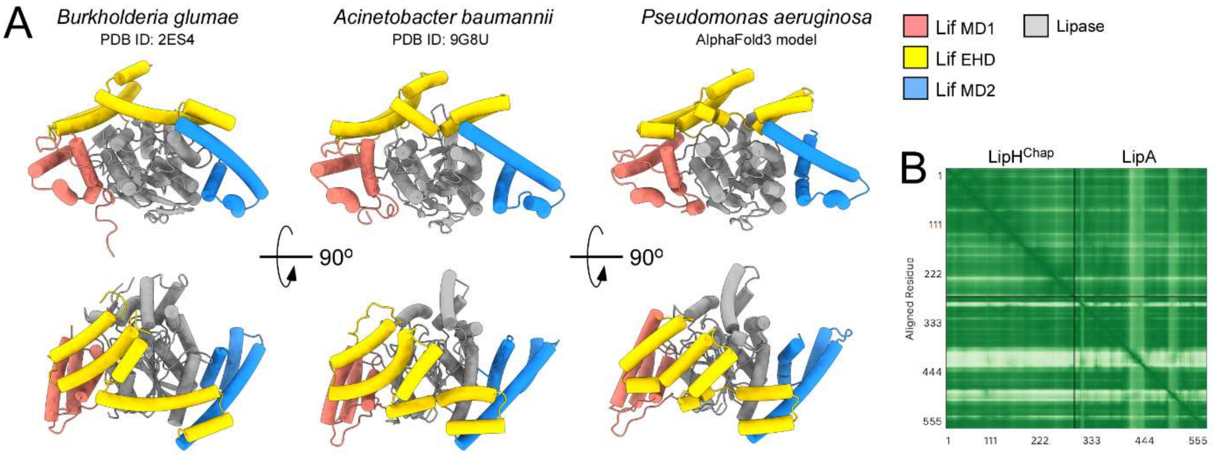


**Supplemental Figure 1. Conserved organization of the foldase:lipase complex.**

**(A)** Crystal structures of lipases from *B. glumae* and *A. baumannii* bound to their cognate foldase chaperones (Lif's), in comparison to the AlphaFold3-derived model for the $LipH^{Chap}$:LipA complex of *P. aeruginosa* (excluding the N-terminal anchor/linker region; ipTM score 0.81).

**(B)** Predicted aligned error plot for the $LipH^{Chap}$:LipA complex of *P. aeruginosa.*

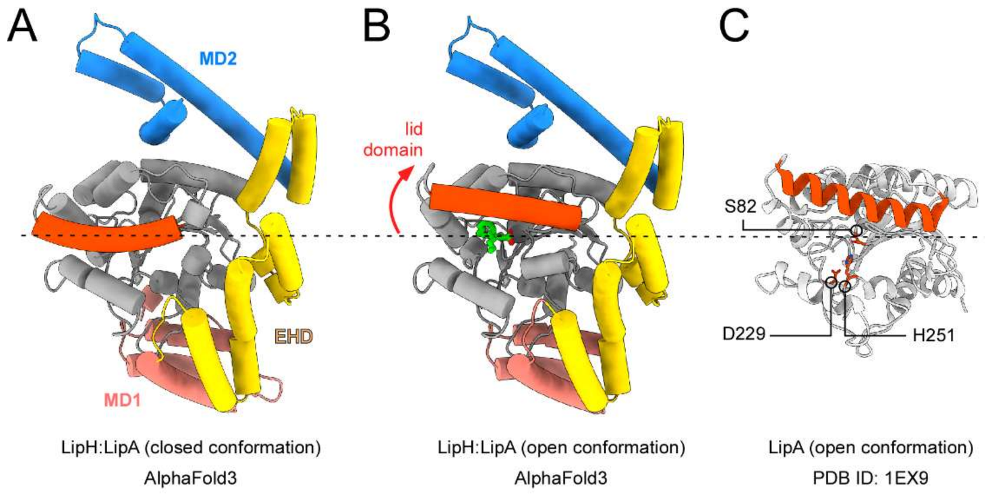


**Supplemental Figure 2. Gating-associated dynamics of *P. aeruginosa* LipA.**

**(A)** AlphaFold3-derived model of the LipH:LipA complex. The gating helix 5 of LipA ("lid domain") is shown in red.

**(B)** AlphaFold3-derived model of the LipH:LipA complex in presence of an oleic acid molecule (green). Docking of the ligand is facilitated by a displacement of the lid domain, as indicated by the arrow.

**(C)** Crystal structure of *P. aeruginosa* LipA shows a displaced lid domain. The residues forming the catalytic triad are indicated.

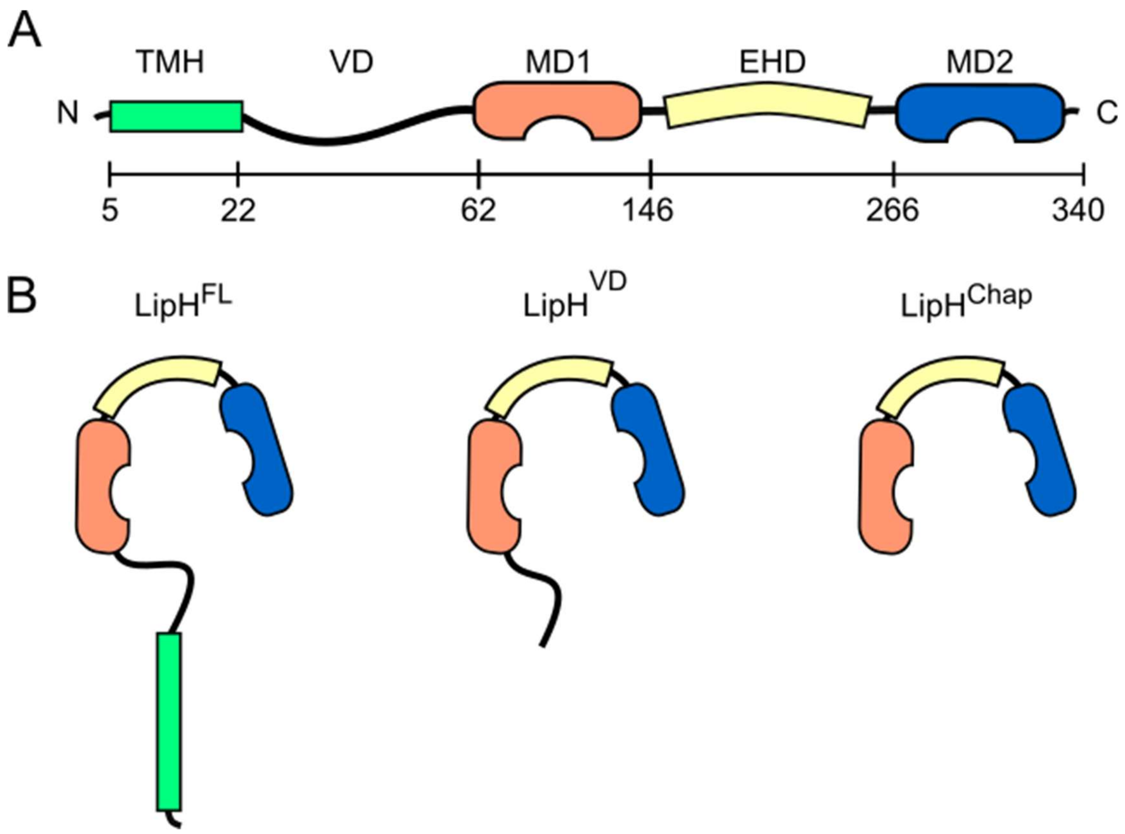


**Supplemental Figure 3. LipH architecture and design of the studied constructs.**

**(A)** Organization of the full-length foldase LipH. The structural elements and their positions within the polypeptide chain (numbers in amino acids) are indicated.

**(B)** Schematic overview of the constructs employed in the study

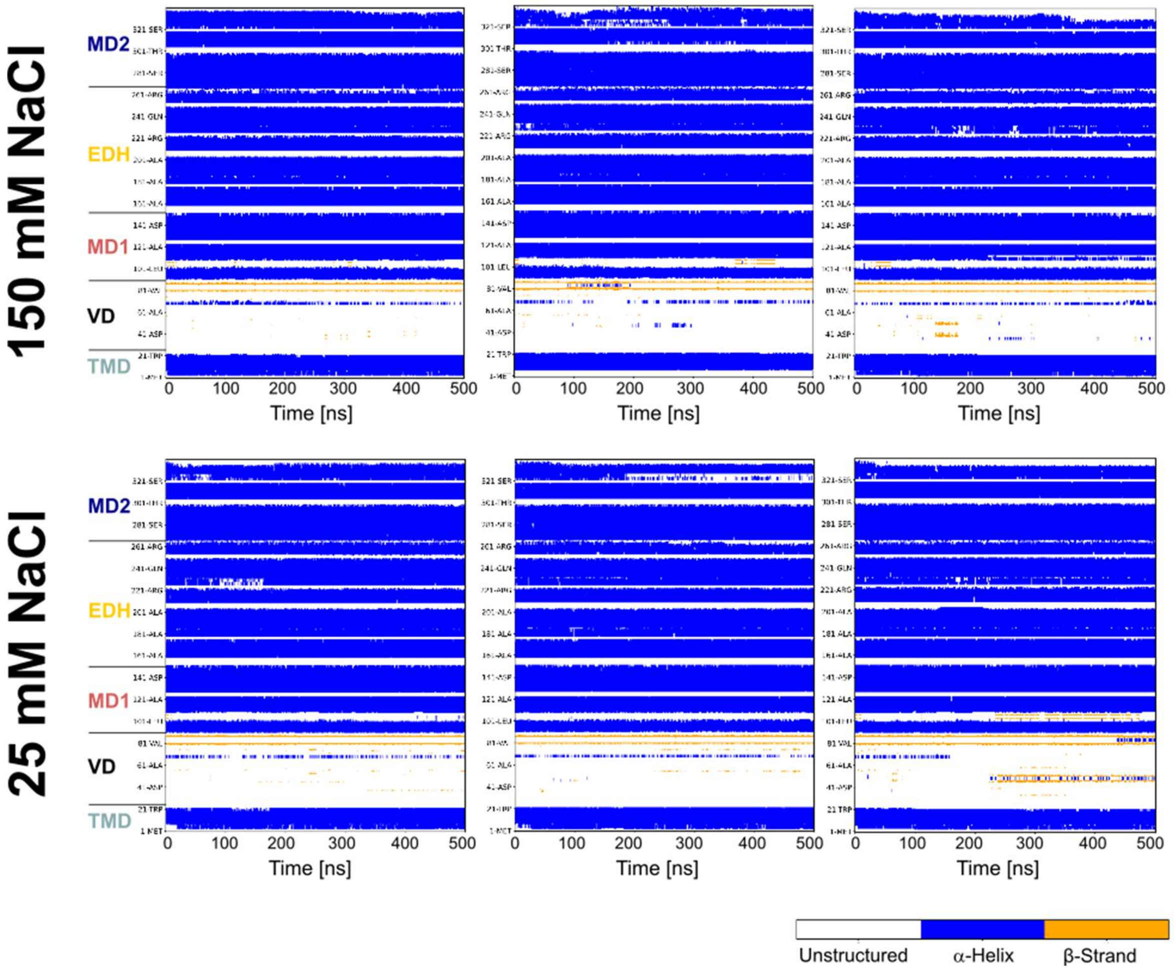


**Supplemental Figure 4. LipH retains its secondary structure over the course of molecular dynamics simulations.**

The color-coded secondary structure of LipH$^{FL}$ along individual simulation courses at 150mM and 25 mM NaCl is plotted against the simulation time. The structural domains of LipH are indicated on the left.

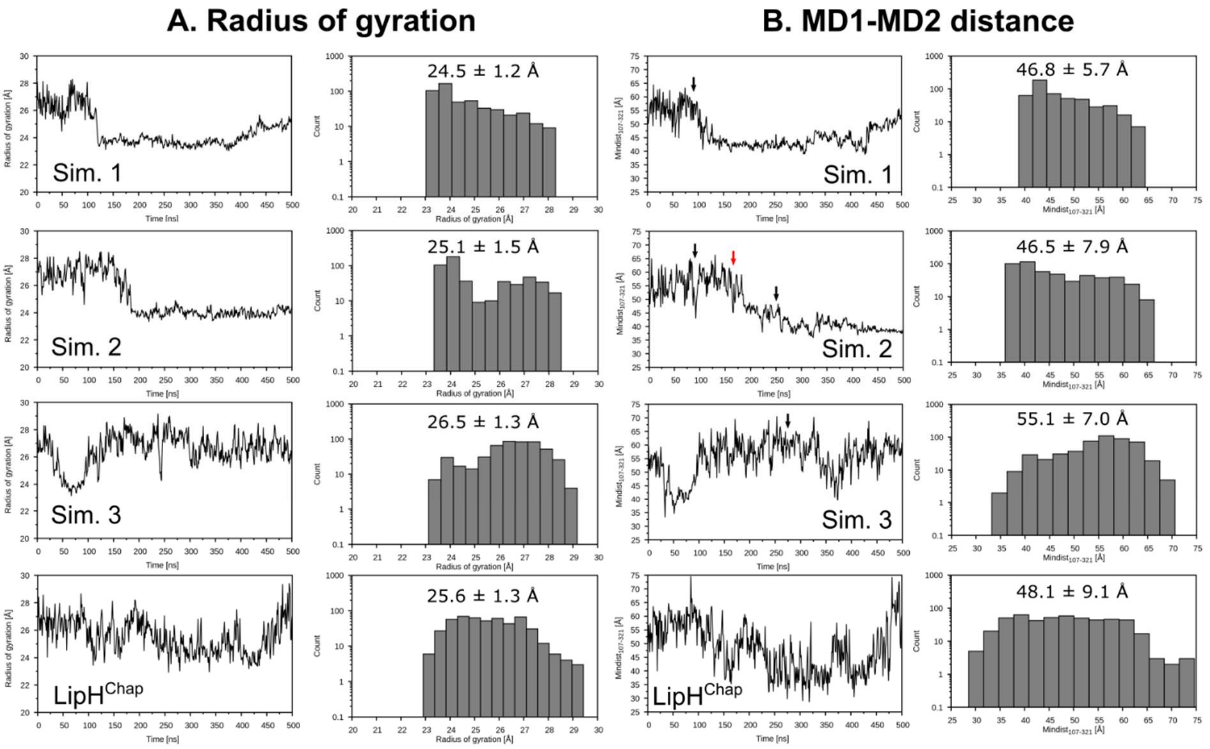


**Supplemental Figure 5. Conformational dynamics of LipH at 150 mM NaCl.**

**(A)** Variations in the radius of gyration of the chaperoning domain within LipH$^{FL}$ over the time course of individual simulations (left) and the corresponding distributions of the radius of gyration (right). The mean value ± standard deviations are indicated for each simulation. "LipH$^{Chap}$" corresponds to the LipH chaperoning domain simulated without the membrane anchor and VD.

**(B)** Variations in the distance between MD1 and MD2 domains within LipH$^{FL}$ over the time course of individual simulations (left) and the corresponding distributions of the distance (right), with the mean value ± standard deviations indicated.

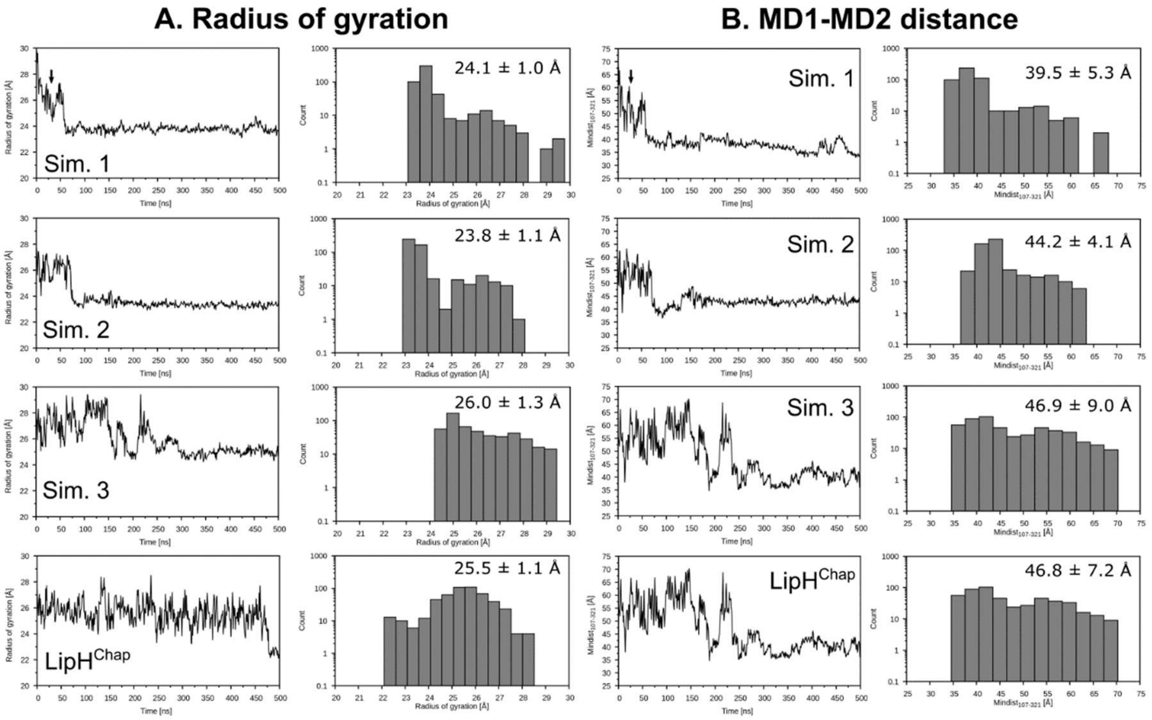


**Supplemental Figure 6. Conformational dynamics of LipH at 25 mM NaCl.**

**(A)** Variations in the radius of gyration of the chaperoning domain within $\text{LipH}^{\text{FL}}$ over the time course of individual simulations (left) and the corresponding distributions of the radius of gyration (right). The mean value ± standard deviations are indicated for each simulation. "$\text{LipH}^{\text{Chap}}$" corresponds to the LipH chaperoning domain simulated without the membrane anchor and VD.

**(B)** Variations in the distance between MD1 and MD2 domains within $\text{LipH}^{\text{FL}}$ over the time course of individual simulations (left) and the corresponding distributions of the distance (right), with the mean value ± standard deviations indicated.

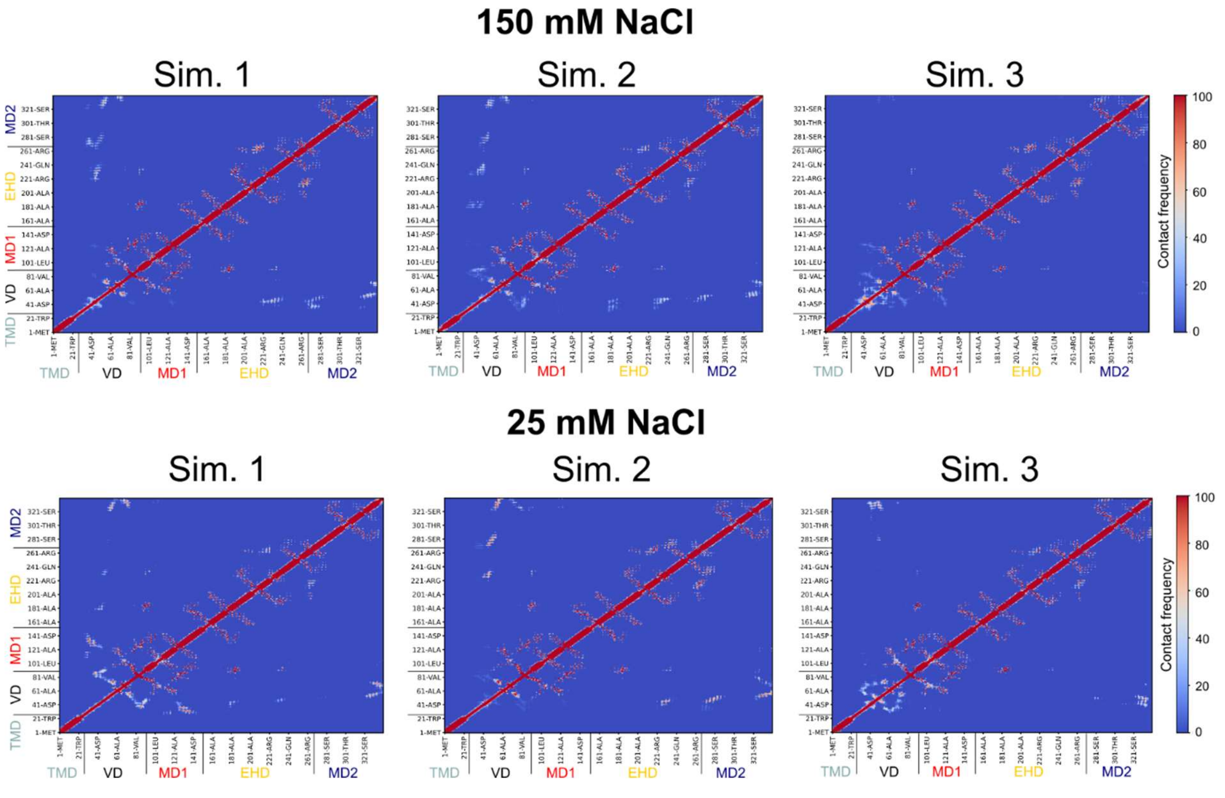


**Supplemental Figure 7.** Maps of intramolecular contacts within LipH$^{FL}$ over the course of individual simulations at 150 mM and 25 mM NaCl. The structural domains of LipH are indicated.

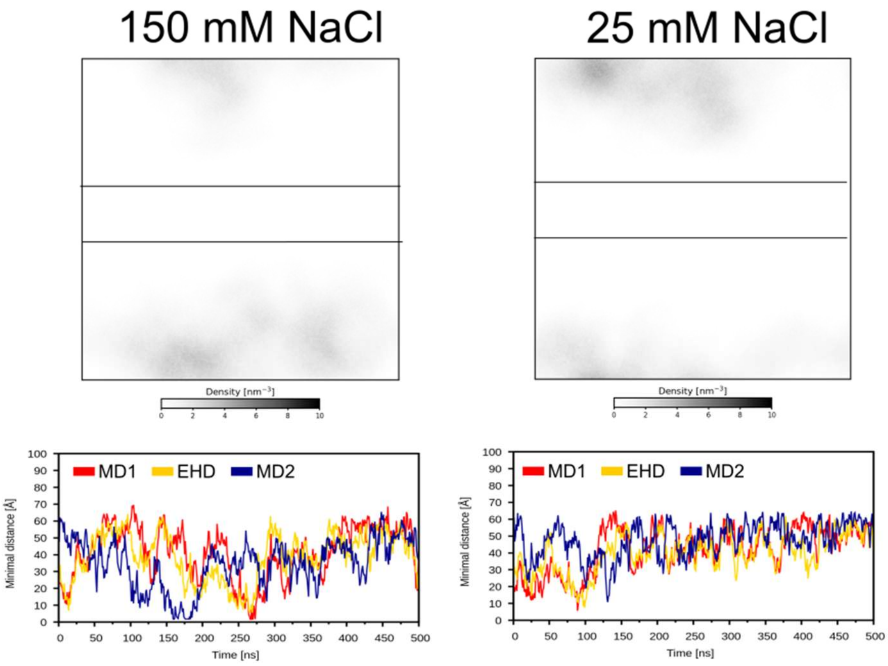


**Supplemental Figure 8. LipH$^{Chap}$ localization along molecular dynamics simulations.**

Top: Density maps of LipH$^{Chap}$ localization within the simulation box calculated for 150 mM and 25 mM NaCl.

Bottom: Corresponding minimal distances of the individual LipH domains to the membrane over the course of each simulation.

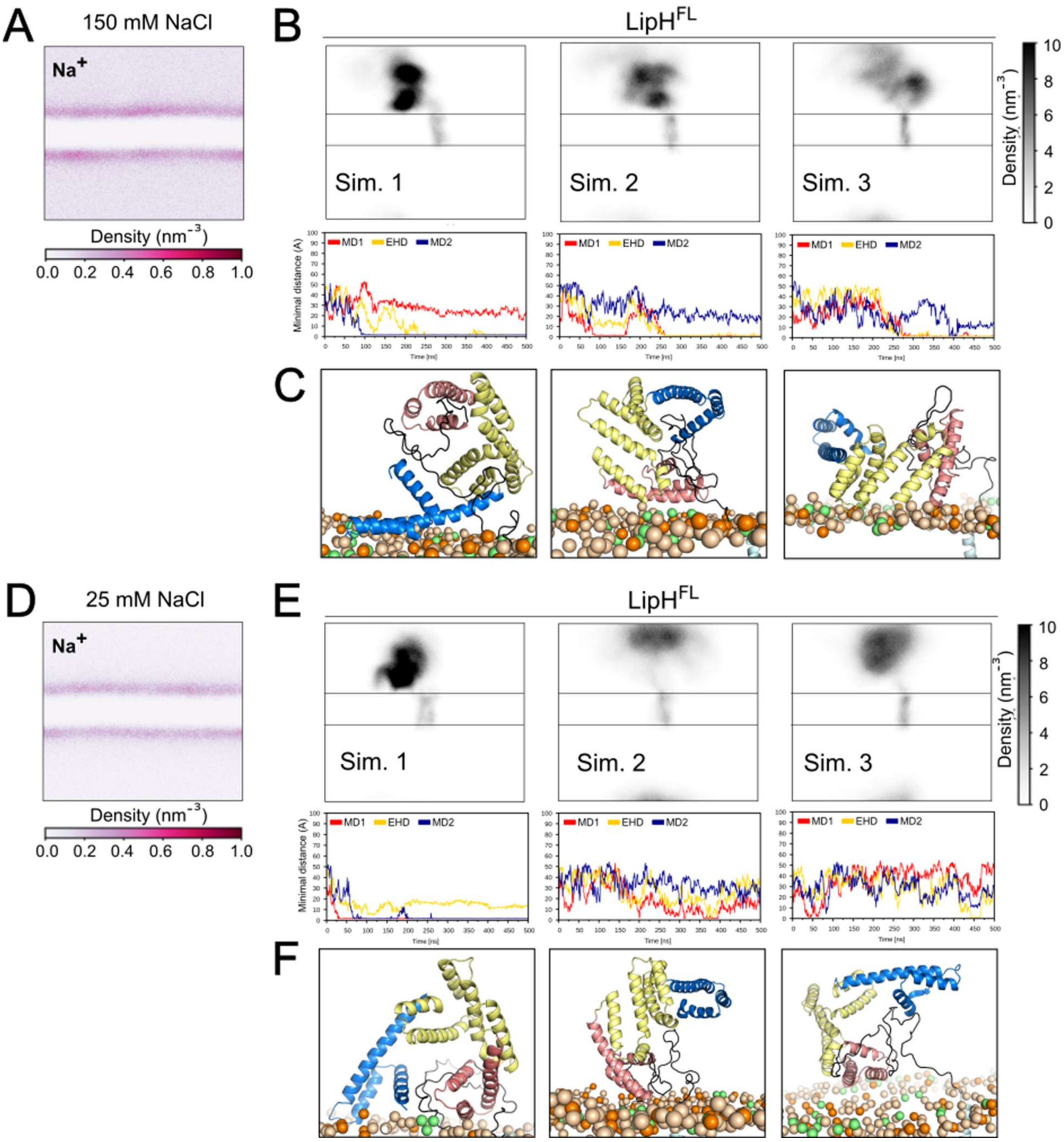


**Supplemental Figure 9. LipH interactions with the membrane interface.**

**(A)** Exemplary density map of sodium ions at 150 mM NaCl (derived from simulation #1) highlights the lipid membrane interface.

**(B)** Density maps of LipH$^{FL}$ localization within the simulation box calculated for individual simulations at 150 mM NaCl. The lipid bilayer borders are indicated as lines. Below: Corresponding minimal distances of the individual LipH domains to the membrane over the course of each simulation.

**(C)** A selection of poses taken by LipH at the membrane interface in simulations at 150 mM NaCl.

**(D-F)** Same as **(A-C)** upon simulations at 25 mM NaCl.

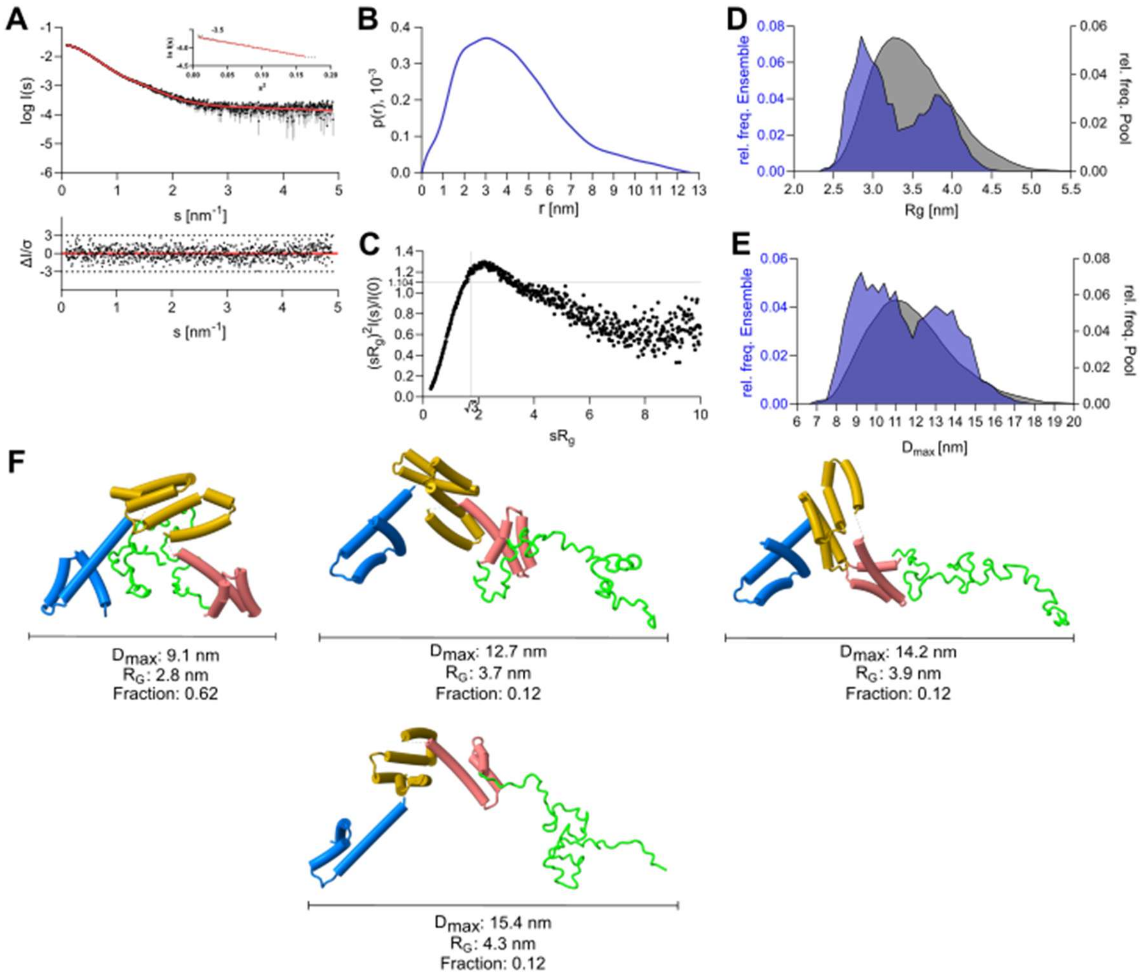


**Supplemental Figure 10. Small-angle X-ray scattering analysis of LipH$^{VD}$.**

**(A)** Experimental data are shown in black dots, with grey error bars. The EOM model fit is shown as red line; below is the residual plot of the data. The Guinier plot of LipH$^{VD}$ is shown in the inset.

**(B)** The pair distance distribution function *p(r)* of LipH$^{VD}$ as determined by SAXS.

**(C)** Dimensionless Kratky plots of LipH$^{VD}$.

**(D)** Relative frequency distribution against the $R_g$ of the EOM model. The selected ensemble is shown in blue, while the relative frequency of the initial pool is shown in grey.

**(E)** Relative frequency distribution of the $D_{max}$ among the EOM model. The selected ensemble is shown in blue, while the relative frequency of the initial pool is shown in grey.

**(F)** Selected EOM models with their corresponding $D_{max}$ and $R_g$ values and the fractions within the ensemble.

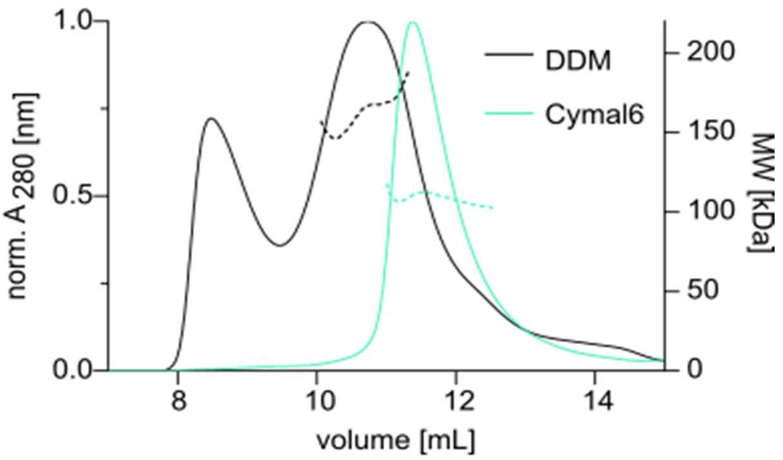


**Supplemental Figure 11. SEC-MALS of the full-length LipH in DDM and Cymal-6 micelles.** Normalized UV absorbance of of LipH$^{FL}$ in DDM (black) and Cymal-6 (petrol) is shown as solid lines. Dashed lines show the molecular weights in kDa determined by MALS (right Y-axis).

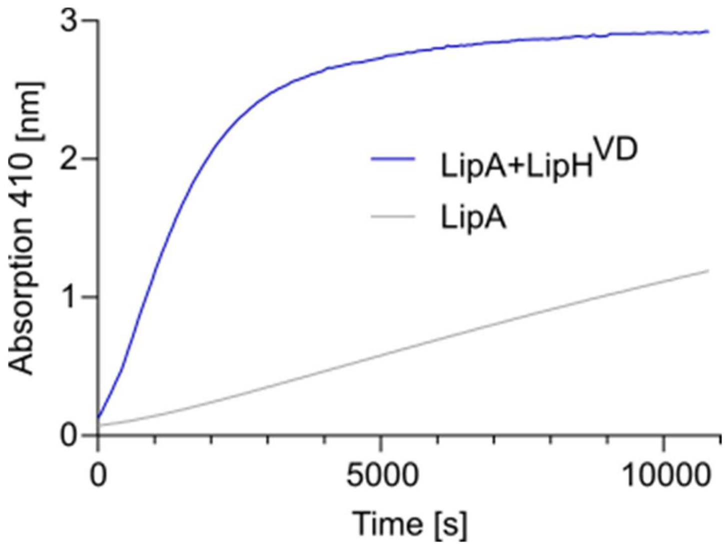


**Supplemental Figure 12: *In vitro* analysis of LipH-mediated enzymatic activity of LipA.**

The lipase activity induced by LipH was monitored via hydrolysis of *p*-nitrophenyl butyrate to *p*-nitrophenolate and butyric acid, measured as an increase in absorbance at 410 nm. The substrate hydrolysis is shown in the presence of the chaperone LipH$^{VD}$ (blue) and in its absence (grey).

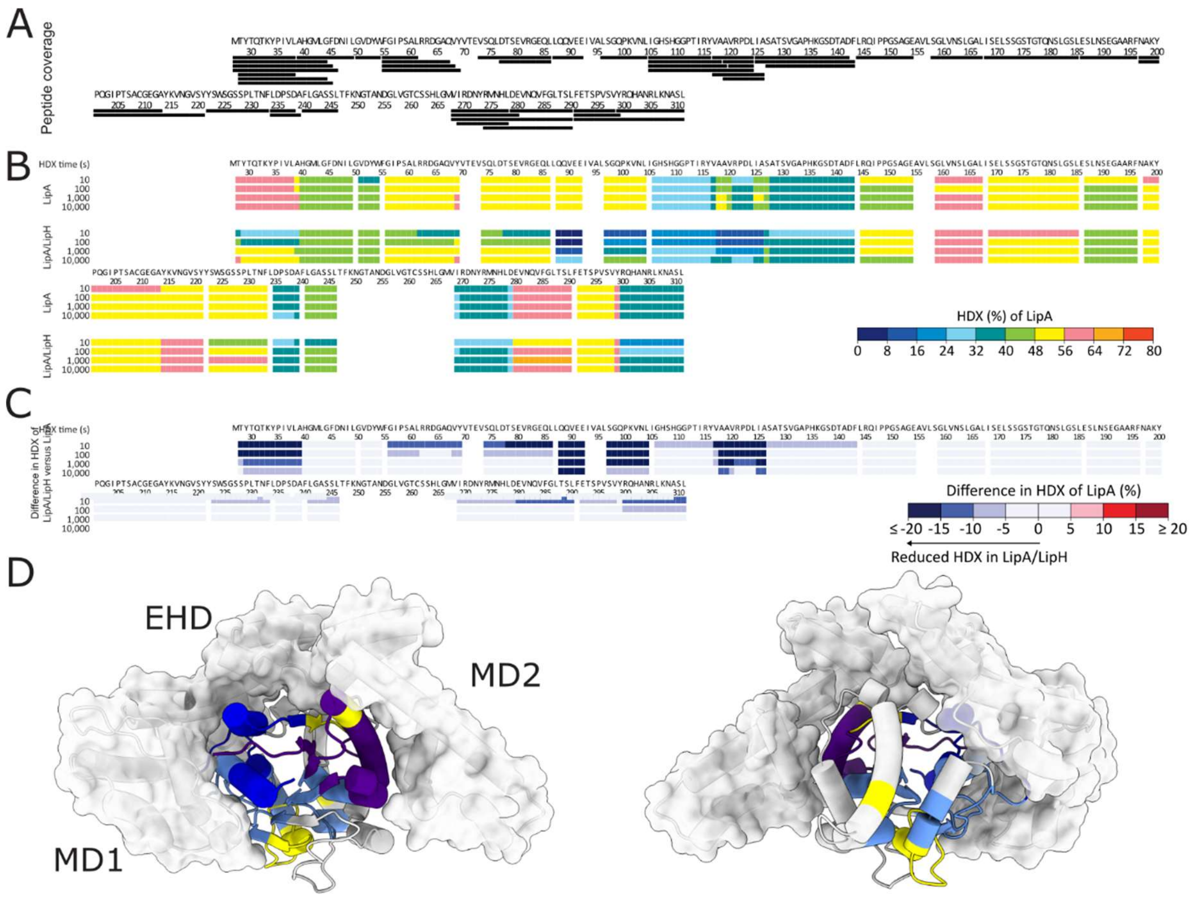


**Supplemental Figure 13. Hydrogen/deuterium exchange mass spectrometry of LipA.**

**(A)** Overview of LipA peptides identified in HDX-MS experiments. Each black bar represents an identified peptide. In total, 49 peptides spanning 89.5% of the LipA$^{F144E}$ amino acid sequence were analyzed for their H/D exchange.

**(B)** The residue-specific HDX of individual LipA and of LipA in presence of LipH$^{Chap}$, color-coded from 0% (blue) to 80% (red).

**(C)** The difference in the residue-specific HDX of LipA in presence of LipH$^{Chap}$ and individual LipA is color-coded from ≤-20% (blue) to ≥20% (red).

**(D)** Differences in the residue-specific HDX levels plotted on LipA structure in complex with LipH$^{Chap}$. The major HDX decrease is observed within the N-terminal fragment of LipA which interacts with MD2 of the chaperone.

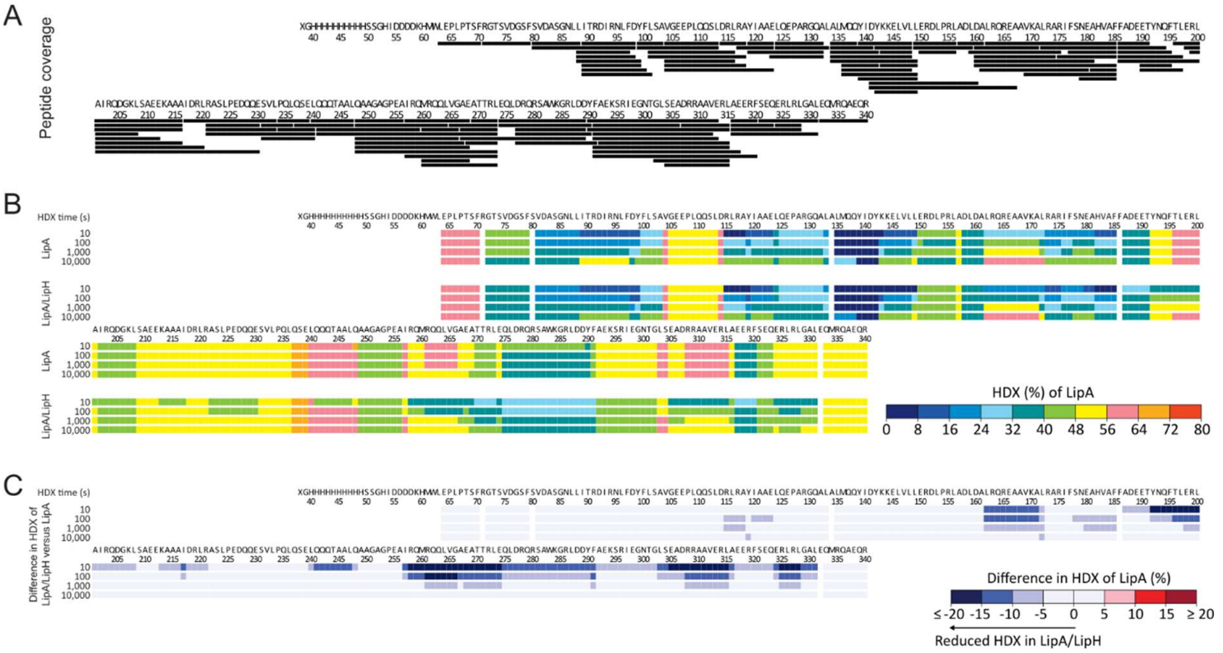


**Supplemental Figure 14. Hydrogen/deuterium exchange mass spectrometry of LipH$^{Chap}$**

**A.** Overview of LipH$^{Chap}$ peptides identified in HDX-MS experiments. Each black bar represents a peptide of LipH$^{Chap}$. In total, 126 peptides spanning 91.7% of the LipH$^{Chap}$ amino acid sequence were analyzed for their H/D exchange.

**B.** The residue-specific HDX of individual LipH$^{Chap}$ and of LipH$^{Chap}$ in presence of LipA is color-coded from 0% (blue) to 80% (red).

**C.** The difference in residue-specific HDX of LipH$^{Chap}$ in presence of LipA and individual LipH$^{Chap}$ is color-coded from ≤-20% (blue) to ≥20% (red).

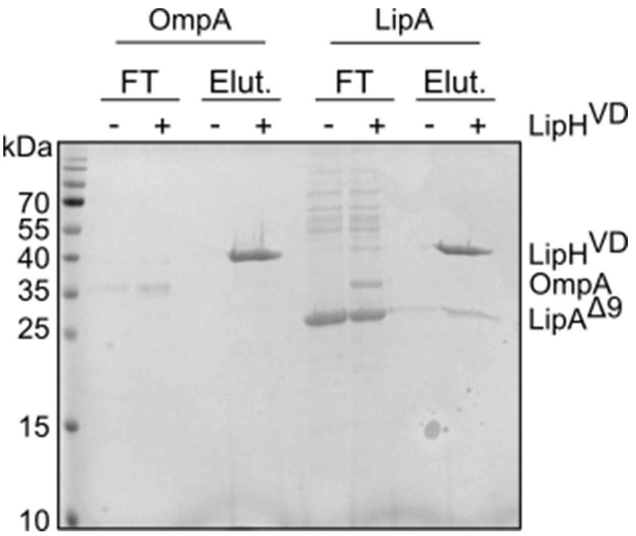


**Supplemental Figure 15. Co-elution interaction assay control with urea-unfolded OmpA.**

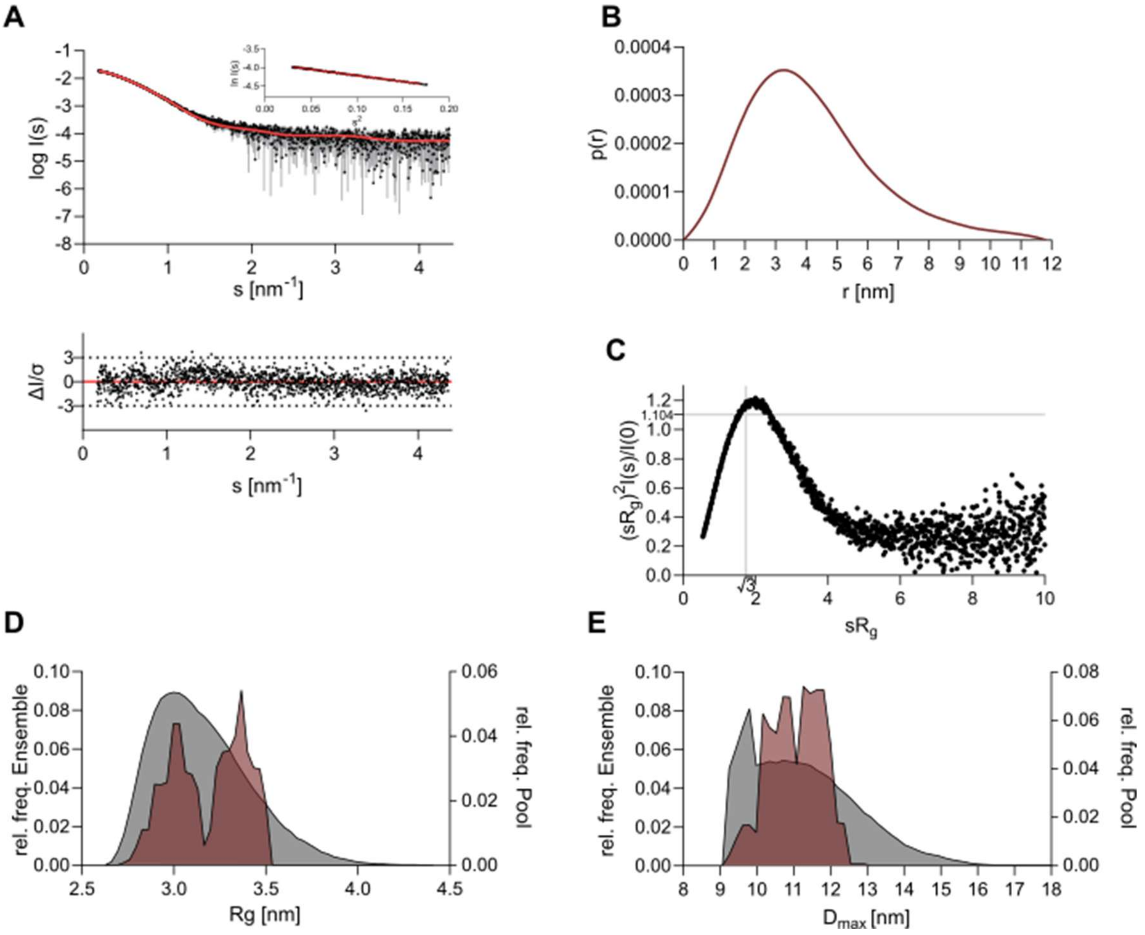


**Supplemental Figure 16. Small-angle X-ray scattering analysis of the LipH$^{VD}$:LipA complex.**

**(A)** Experimental data are shown in black dots, with grey error bars. The EOM ensemble fit is shown as red line; below is the residual plot of the data. The Guinier plot of LipH$^{VD}$:LipA is shown in the inset.

**(B)** The pair distance distribution function p(r) of LipH$^{VD}$:LipA complex as determined by SAXS.

**(C)** Dimensionless Kratky plots of LipH$^{VD}$:LipA complex.

**(D)** Relative frequency distribution against the $R_g$ of the EOM model. The ensemble is shown in brown, while the relative frequency of the pool is shown in grey.

**(E)** Relative frequency distribution against the $D_{max}$ of the EOM model. The ensemble is shown in brown, while the relative frequency of the pool is shown in grey.

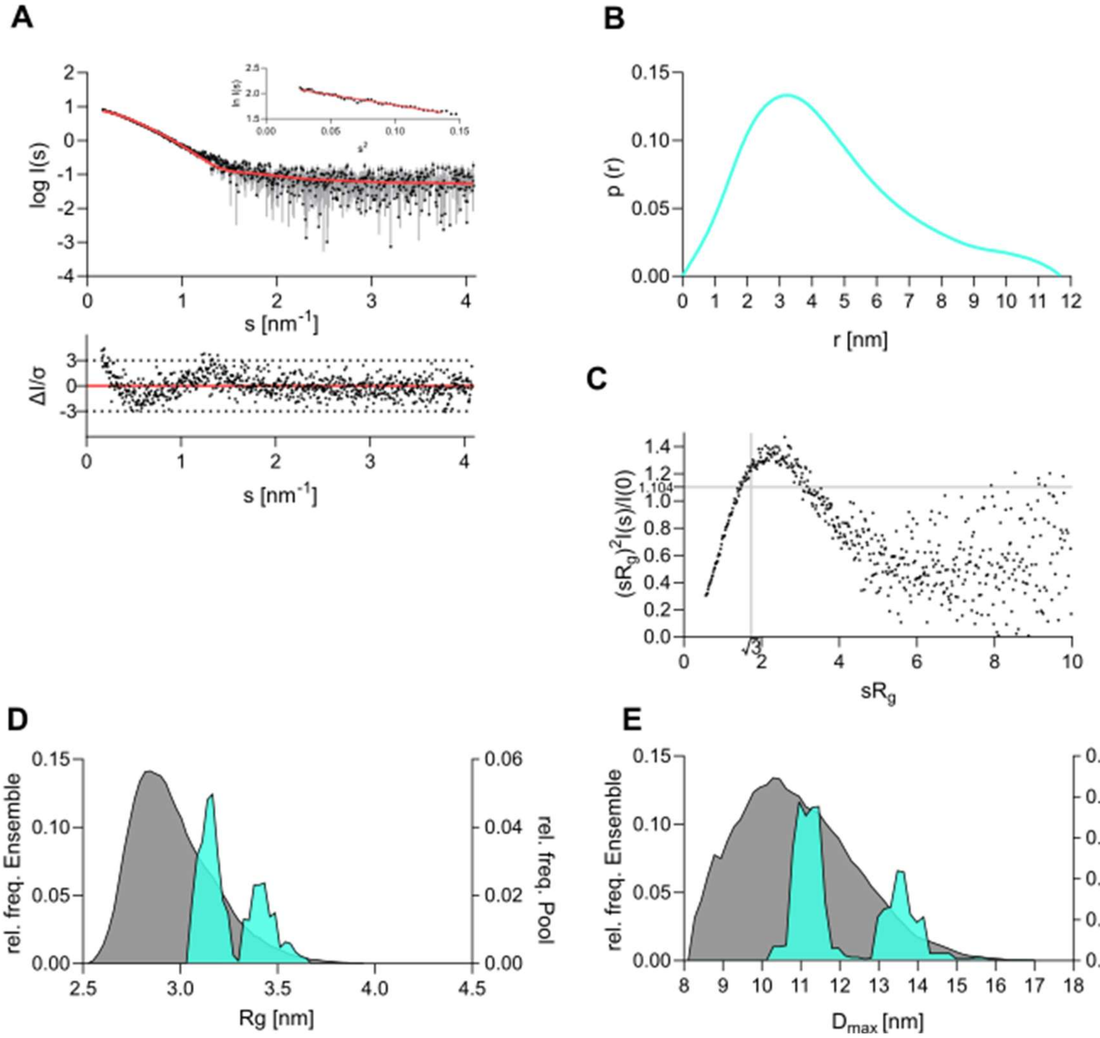


**Supplemental Figure 17. Small-angle X-ray scattering analysis of the LipH$^{VD}$:LipA$^{\Delta 81}$ complex**

**(A)** Experimental data are shown in black dots, with grey error bars. The EOM ensemble fit is shown as red line; below is the residual plot of the data. The Guinier plot of LipH$^{VD}$:LipA$^{\Delta 81}$ is shown in the inset.

**(B)** The pair distance distribution function *p(r)* of LipH$^{VD}$:LipA$^{\Delta 81}$ complex as determined by SAXS.

**(C)** Dimensionless Kratky plots of LipH$^{VD}$:LipA$^{\Delta 81}$ complex.

**(D)** Relative frequency distribution against the $R_g$ in nm of the EOM model. The ensemble is shown in cyan, while the relative frequency of the pool is shown in grey.

**(E)** Relative frequency distribution against the $D_{max}$ in nm of the EOM model. The ensemble is shown in cyan, while the relative frequency of the pool is shown in grey.

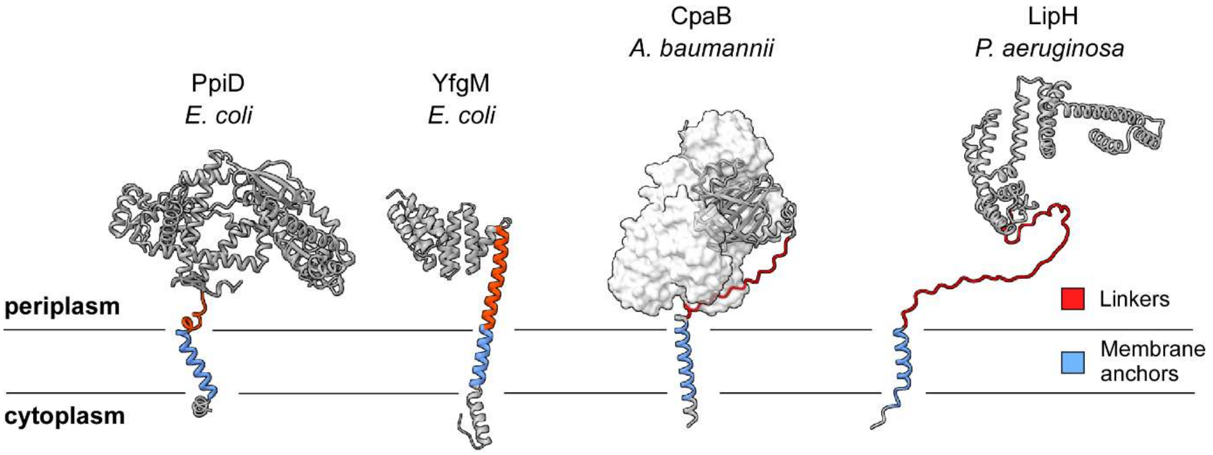


**Supplemental Figure 18. AlphaFod3-based models of the membrane-anchored chaperones.** Membrane anchors are predicted based on the hydrophobicity via TMHMM. For *A. baumannii* CpaB chaperone, its cognate client protease CpaB is shown (surface visualization).

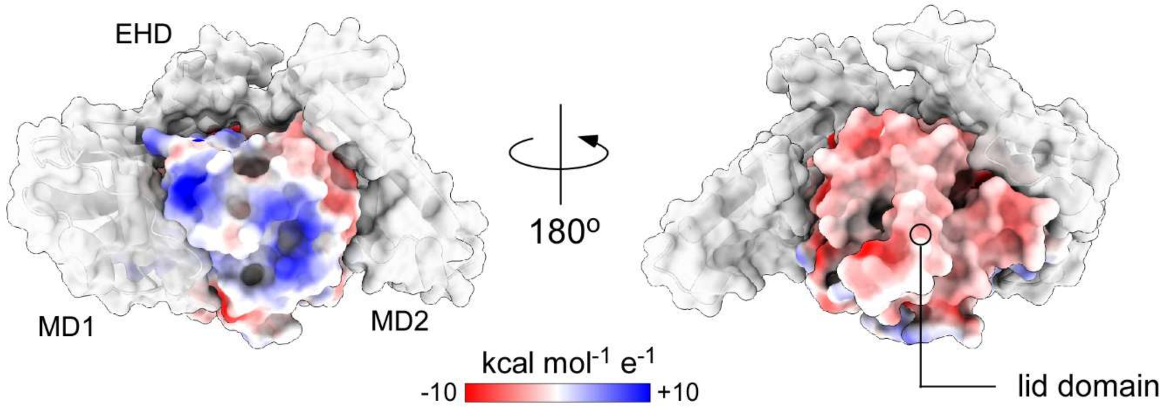


**Supplemental Figure 19. The electrostatic potential at the surface of the folded LipA.**

The molecular surface of LipA is coloured according to the local electrostatic potentials (blue = cationic; red = anionic; the scale bar shown below). The chaperoning domain of LipH is shown as a semi-transparent grey surface, with positions of the structural sub-domains indicated.

**Supplemental Table 1. Summary of SAXS data acquisition and analysis**

| | | | |
|---|---|---|---|
| **Data collection parameters** | | | |
| SAXS Device | Xenocs Xeuss 2.0 with Q-Xoom | P12, PETRA III, DESY Hamburg (Blanchet et al. 2015) | BM29, ESRF Grenoble (Tully et al. 2023) |
| Detector | PILATUS 3 R 300K windowless | PILATUS 6 M | PILATUS3 x 2 M |
| Detector distance (m) | 0.550 | 3.0 | 2.827 |
| Beam size | 0.8 mm x 0.8 mm | 120 µm x 200 µm | 200 µm x 100 µm |
| Wavelength (nm) | 0.154 | 0.124 | 0.099 |
| Sample environment | Low Noise Flow Cell, 1 mm ø | Quartz glass capillary, 1 mm ø | |
| Absolute scaling method | Comparison with scattering from pure $H_2O$ | | |
| Normalization | To transmitted intensity by beam-stop counter or direct beam | | |
| Scattering intensity scale | Absolute scale, $cm^{-1}$ | | |
| *s* range ($nm^{-1}$)‡ | 0.05 – 5.5 | 0.03 – 7.0 | 0.025–5.5 |
| **Sample** | **$LipH^{VD}$** | **$LipH^{VD}$:LipA complex** | **$LipH^{VD}$:$LipA^{\Delta 81}$ complex** |
| Organism | *Pseudomonas aeruginosa PAO1* | | |
| UniProt ID | Q01725 | LipH: Q01725<br>LipA: P26876 | LipH: Q01725<br>LipA: P26876 |
| Mode of measurement | batch | Online SEC-SAXS | |
| SEC-Column | - | Superdex 200 Increase 10/300 GL | |
| Flowrate (mL/min) | - | 0.6 | 0.6 |
| Injection volume (µL) | - | 100 | 100 |
| Temperature (°C) | 10 | 20 | 20 |
| Exposure time s (# frames) | 600 (24) | 0.995 (2400) | 2 (1200) |
| # frames used for averaging | 20 | 24 | 18 |
| Protein buffer | 50 mM Tris, 100 mM NaCl, 100 µM TCEP, 5% glycerol, pH 8.0 | 5 mM Tris, 5 mM glycine, 1 mM $CaCl_2$, 5% glycerol, pH 8.0 | 5 mM Tris, 5 mM glycine, 1 mM $CaCl_2$, 5% glycerol, pH 8.0 |
| Protein concentration [mg/mL] | 11.82 | 8 mg/mL $LipH^{VD}$ + 0.8 mg/mL $LipA^{FL}$ | 1 mg/mL $LipH^{VD}$ + $LipA^{d81}$ |
| **Structural parameters** | | | |
| *Guinier Analysis (PRIMUS)* | | | |
| *I*(0) ± s ($cm^{-1}$) | 0.025 ± 0.0001 | 0.021 ± 0.00004 | 8.834 ± 0.072 |
| $R_g$ ± s (nm) | 3.22 ± 0.022 | 3.14 ± 0.001 | 3.51 ± 0.043 |
| *s-range* ($nm^{-1}$) | 0.088 – 0.403 | 0.173 – 0.411 | 0.163 – 0.369 |
| *min < sRg < max limit* | 0.282 – 1.297 | 0.544 – 1.289 | 0.572 – 1.294 |
| Data point range | 1 – 55 | 1 – 85 | 1 – 43 |
| Linear fit assessment ($R^2$) | 0.990 | 0.996 | 0.972 |

| |
|---|
| *PDDF/P® Analysis (GNOM)* |

| | | | |
|---|---|---|---|
| $I(0) \pm s$ ($cm^{-1}$) | 0.026 ±0.0001 | 0.021 ± 0.00003 | 8.874 ± 0.058 |
| $R_g \pm s$ (nm) | 3.41 ± 0.026 | 3.26 ± 0.0095 | 3.53 ± 0.028 |
| $r_{max}$ (nm) | 12.61 | 11.82 | 11.70 |
| Porod volume ($nm^3$) | 69.97 | 110.09 | 103.55 |
| *s-range* ($nm^{-1}$) | 0.088 – 4.895 | 0.173 – 4.360 | 0.163 – 4.081 |
| χ2 / CorMap P-value | 1.015 / 0.553 | 1.003 / 0.087 | 1.069 / 0.092 |
| **Molecular mass (kDa)** | | | |
| From *I*(0) | 34.62 | n.d. | n.d. |
| From Qp (Porod 1951) | 40.03 | 69.34 | 63.04 |
| From MoW2 (Fischer et al. 2010) | 34.20 | 68.07 | 51.40 |
| From Vc (Rambo and Tainer 2013) | 38.87 | 66.58 | 55.40 |
| Bayesian Inference (Hajizadeh et al. 2018) | 37.70 | 67.08 | 58.15 |
| From sequence | 38.32 | 68.46 (1:1 stoichiometry) | 59.61 (1:1 stoichiometry) |
| **Flexibility ensemble modelling** | | | |
| EOM | | | |
| Symmetry | P1 | P1 | P1 |
| *s*-range for fit ($nm^{-1}$) | 0.088 – 4.895 | 0.173 – 4.360 | 0.163 – 4.081 |
| $\chi^2$, CorMap *P*-value | 1.026 / 0.553 | 1.136 / 0.166 | 1.528 / 0. 000003 |
| **SASBDB accession codes** (Kikhney et al. 2020) | xx | xx | xx |
| **Software** | | | |
| ATSAS Software Version (Manalastas-Cantos et al. 2021) | 3.0.5 | | |
| Primary data reduction | CHROMIXS (Panjkovich and Svergun 2018)/ PRIMUS (Konarev et al. 2003) | | |
| Data processing | GNOM (Svergun 1992) | | |
| Flexibility ensemble modellin | EOM (Bernado et al. 2007; Tria et al. 2015) | | |
| Statistic goodness-of-fit test | χ 2, CorMap (Franke, Jeffries, and Svergun 2015) | | |
| Model visualization | Chimera X 1.10 (Goddard et al. 2018) | | |

‡s = 4πsin(θ)/λ, 2θ – scattering angle, n.d. not determined
